# Loss of FMRP in microglia promotes degeneration of parvalbumin neurons and audiogenic seizures via progranulin insufficiency

**DOI:** 10.1101/2025.07.09.663876

**Authors:** Heng Li, Jennifer Ja-Yoon Choi, Eric J. Huang, Baoji Xu

**Author notes:** Correspondence: Dr. Baoji Xu, Department of Neuroscience, The Herbert Wertheim UF Scripps Institute for Biomedical Innovation & Technology, University of Florida, 130 Scripps Way, Jupiter, FL 33458, USA.

## Abstract

Fragile X syndrome (FXS) results from loss of FMR1-encoded FMRP and is associated with reduced density of parvalbumin (PV) neurons; however, the mechanism underlying this abnormality remains unknown. Here we report that microglial FMRP regulates PV neuron density through lysosomal function. Mice with *Fmr1* deletion in microglia exhibited audiogenic seizures (AGS) and decreased PV neuron density in the cortex and AGS-associated inferior colliculus (IC). FMRP increased the expression of lysosomal genes in microglia, including the progranulin-encoding *Grn* gene. Its loss in microglia led to impaired lysosomal function and increased apoptosis in microglia and PV neurons. Furthermore, PV neuron density in the IC was reduced similarly in male *Grn*^+/-^, *Fmr1*^-/y^, and *Grn*^+/-^;*Fmr1*^-/y^ mice, and AAV8-mediated overexpression of progranulin rescued AGS and PV neuron loss in *Fmr1*^-/y^ mice. This indicates that progranulin insufficiency is a determinant for PV neuron loss in FXS and elevating progranulin is a therapeutic strategy for FXS.

---

Fragile X syndrome (FXS) is the leading cause for inheritable intellectual disability and monogenic autism spectrum disorder (ASD)^1–3^. Individuals with FXS display a range of cognitive and behavioral deficits, including anxiety, hyperactivity, memory deficits, and seizures^4,5^, and ∼50% of males with FXS meet the diagnostic criteria for ASD^6,7^. Currently, there is no approved drug for FXS treatment. FXS results from transcriptional silencing of the *FMR1* gene which encodes the RNA-binding fragile X messenger ribonucleoprotein (FMRP)^8,9^. Nearly all cases of FXS are caused by the trinucleotides CGG repeats expansion in the 5′ untranslated region of the *FMR1* gene, resulting in hypermethylation and absence of FMRP^8,9^. FMRP binds hundreds of mRNAs in the brain and is involved in many aspects of mRNA metabolism^10,11^. In general, FMRP suppresses translation of target mRNAs^12–15^, largely by stalling ribosomal translocation via interacting with the coding region^11^. Additional roles of FMRP in mRNA metabolism include inhibition^16,17^ or activation^18,19^ of translation initiation, regulation of mRNA stability^20,21^, and mRNA editing^22,23^. The protein products of these mRNA targets regulate several biological processes, including axonal and dendritic development, synaptic development and function, cytoskeleton organization, cellular signaling, RNA transport, and transcriptional and epigenetic regulations^3^.

Recent studies have discovered reduced density of PV neurons in multiple cortical regions of FXS patients and mouse models^24–27^. Reduced density or hypoactivity of PV neurons has also been observed in postmortem tissue from individuals with idiopathic ASD^28,29^ and several ASD mouse models including mouse models for sporadic ASD^30–33^, Angelman syndrome^34^, and Rett syndrome^35^. In fact, this PV neuron abnormality in the cortex has become one of the most exciting discoveries in ASD due to incredible convergence between human and animal studies^24^. Given that individual PV neurons in the cortex can innervate and inhibit more than 1,000 excitatory pyramidal neurons^36^, their loss or hypoactivity would have dramatic consequences on network excitability and function and could be a fundamental cause for FXS and other psychiatric disorders^24,37^. However, the mechanism underlying this PV neuron abnormality is unknown.

Microglia are the resident immune cells of the CNS with many fine and motile processes that continuously survey the parenchymal environment to remove cell debris, protein aggregates and pathogens^38,39^. Furthermore, microglia regulate the development of central neural circuits by removing excessive immature neurons^40–43^ and engulfing weak synapses^44–47^. Despite of microglial roles in neuronal development, it remains to be determined whether loss of FMRP expression in microglia contributes to FXS, an X-linked neurodevelopmental disorder. Here, we report that FMRP supports survival of PV neurons by maintaining the expression of lysosomal proteins in microglia during the postnatal brain development. The affected lysosomal proteins include progranulin, which is largely expressed in microglia and regulates lysosomal function and whose haploinsufficiency promotes neurodegeneration^48–52^. Excitingly, our results further show that overexpressing progranulin in the brain immediately after birth prevents loss of PV neurons and audiogenic seizures (AGS) in *Fmr1* null mice, a mouse model for FXS.

## Results

### Selective deletion of the *Fmr1* gene in microglia leads to audiogenic seizures

The role of FMRP in microglia has not been determined yet. To examine the expression of FMRP in microglia, we performed fluorescent immunohistochemistry with antibodies against FMRP and a microglial marker Iba1 on brain sections from C57BL/6J mice of various ages. We found that FMRP was expressed in microglia in all brain regions we examined during late embryogenesis and postnatal brain development but the expression decreased with age and was nearly undetectable by approximately 10 weeks old (Fig. 1a-d and Extended Data Fig. 1a-d). The expression of FMRP in microglia of young mice was also previously reported^53^.

**Fig. 1:**
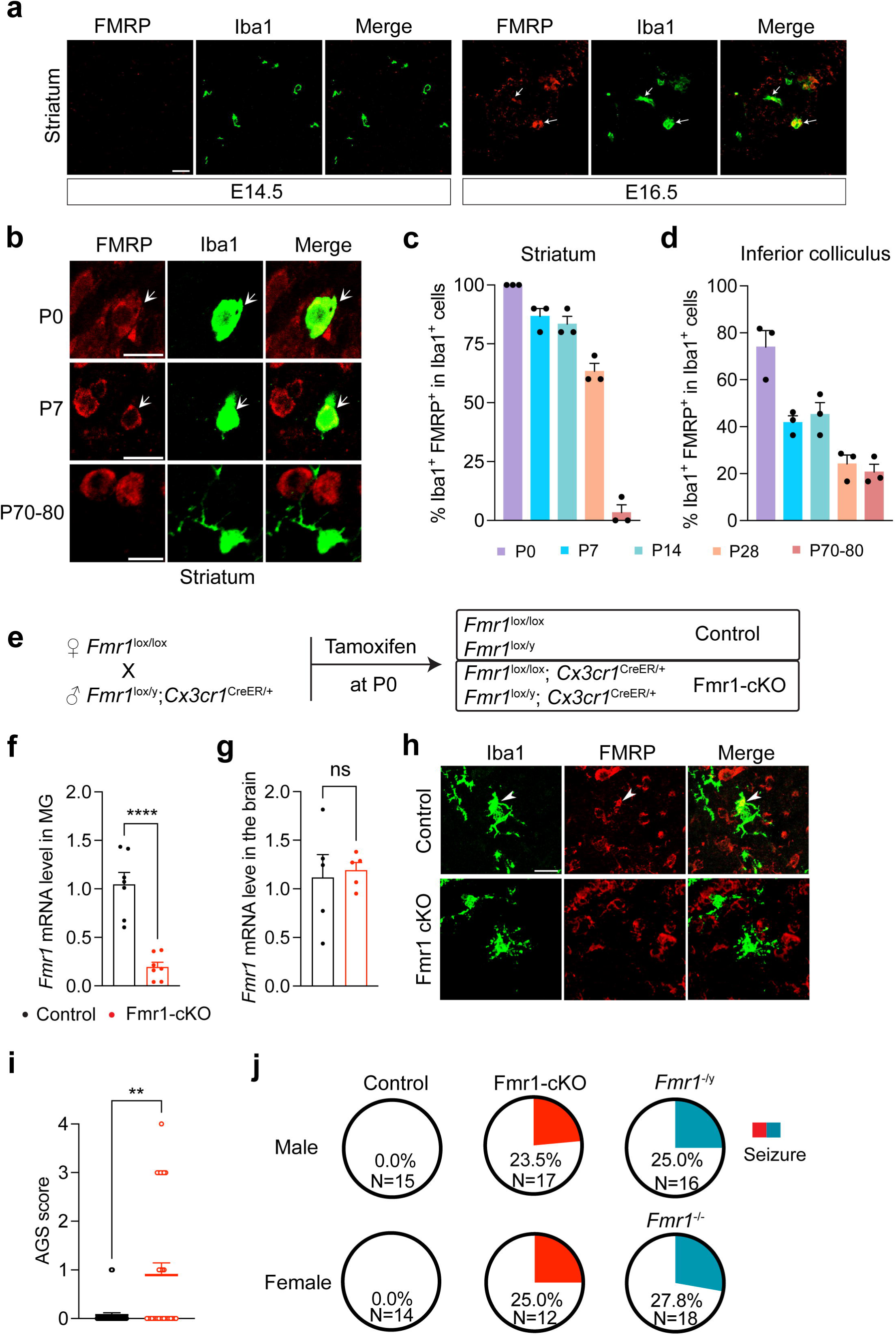
Conditional knockout of the *Fmr1* gene in microglia leads to audiogenic seizures. **a**, Confocal images showing coexpression of FMRP and Iba1 in the striatum of mouse embryonic brains. White arrows indicate colocalization of FMRP and Iba1. Scale bar, 20 μm. **b**, Representative confocal images showing coexpression of FMRP and Iba1 in the striatum of mouse brains at various postnatal ages. White arrows indicate colocalization of FMRP and Iba1. Scale bars, 10 μm. **c** and **d**, Percentage of microglia with FMRP expression in the striatum (**c**) and inferior colliculus (**d**) in mice at P0, P7, P14, P28, and P70-80. n=3 mice of both sexes for each age. **e**, Schematic of the strategy for conditional *Fmr1* deletion in microglia. **f**, The *Fmr1* mRNA level was significantly decreased in microglia purified from Fmr1-cKO mice at P7. Unpaired t test, ****p<0.0001, n=7 mice of both sexes per genotype. **g**, The *Fmr1* mRNA level was not changed in whole-brain lysates of Fmr1-cKO mice at P7. Unpaired t test, ns: not significant, n=5 mice of both sexes per genotype. **h**, Immunohistochemistry showing that FMRP was present in microglia of P14 control mice but was absent in microglia of P14 cKO mice. Images were taken from the hippocampus. Arrows indicate colocalization. Scale bar, 20 μm. **i**, AGS scores in control and Fmr1-cKO mice. Mann-Whitney test, **p=0.0027, n=29 mice for each genotype, including both males and females. **j**, Pie graphs showing percentage of mice developed seizure (AGS score >=2).

Given the expression of FMRP in microglia of the developing brain, we reasoned that loss of FMRP in microglia could alter brain development and contribute to the pathogenesis of FXS. To test this hypothesis, we generated microglia-specific *Fmr1* conditional knockout (Fmr1-cKO) and control mice by employing a floxed *Fmr1* (*Fmr1^lox^*) mouse strain^54^ and an inducible *Cx3cr1^CreER/+^*Cre driver strain^55^ in combination with a single dose of subcutaneous tamoxifen injection at postnatal day 0 (P0; Fig. 1e). The *Cx3cr1^CreER^*driver specifically induced tdTomato expression from the Ai9 Cre reporter^56^ in nearly all microglia after a single dose of tamoxifen injection at P0 (Extended Data Fig. 2a-b). Consistent with this high specificity and efficiency, the *Fmr1* mRNA level was drastically decreased in microglia isolated from Fmr1-cKO mice at P7 compared to control microglia (Fig. 1f and Extended Data Fig. 2c), while remaining unaltered in RNA samples prepared from whole brains of Fmr1-cKO mice (Fig. 1g). Furthermore, FMRP immunoreactivity was abolished in microglia of Fmr1-cKO mice at P14 (Fig. 1h). These results validate selective deletion of the *Fmr1* gene in microglia of Fmr1-cKO mice.

AGS is one of the most robust and consistent behavioral abnormalities observed in *Fmr1* null mice^57,58^. This behavior reveals seizure and hypersensitivity to auditory stimuli of the affected mice. We thus examined AGS in Fmr1-cKO mice at P22-P25. Fmr1-cKO mice were more susceptible to AGS than control mice (Fig. 1i-j). The susceptibility was not due to the mutation in the *Cx3cr1* gene, as *Cx3cr1^CreER/+^* mice and wild-type (WT) littermates had comparable susceptibility to AGS (Extended Data Fig. 2d). Furthermore, the penetrance of AGS was comparable between Fmr1-cKO mice and *Fmr1* null (*Fmr1^-/-^* or *Fmr1*^-/y^) mice (Fig. 1j). These results indicate that the loss of microglial FMRP accounts for the susceptibility to AGS in *Fmr1* null mice.

### *Fmr1* deletion alters microglial morphology

As dysfunction of the IC is implicated as the cause for increased susceptibility to AGS in *Fmr1* null mice^59^, we focused subsequent studies on molecular and cellular alterations in the IC of Fmr1-cKO mice. Microglia in the IC were larger in both sexes and denser in male Fmr1-cKO mice at P14 than age- and sex-matched control mice (Fig. 2a-b and Extended Data Fig. 3a). Microglia in the hippocampus and striatum were also larger in male Fmr1-cKO mice than those in control mice at P14, although their size in females and their density in both sexes were comparable between Fmr1-cKO and control mice (Extended Data Fig. 3b-c). This is similar to the previous observation made in the hippocampus and striatum of *Fmr1* null mice^60^.

**Fig. 2:**
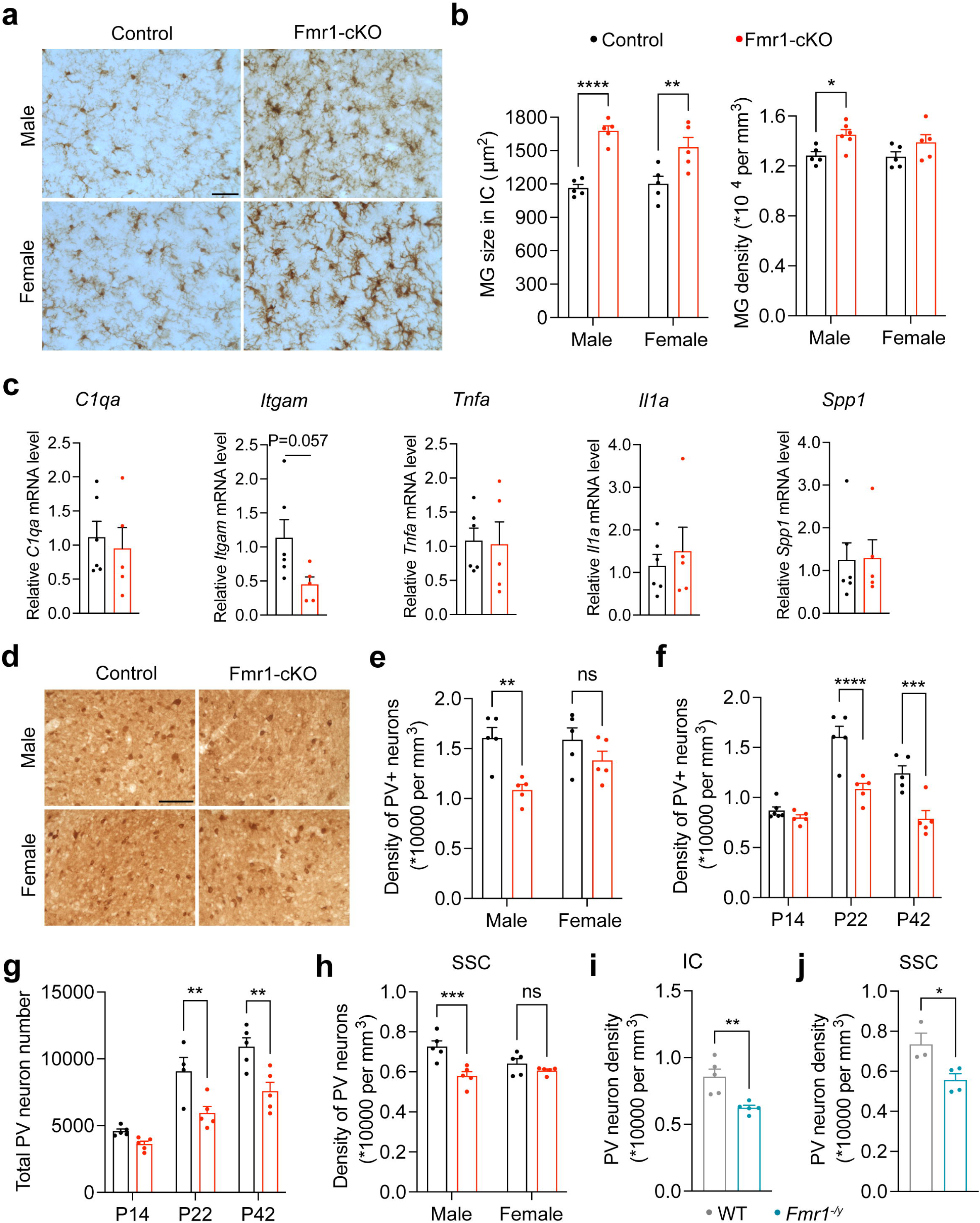
PV neuron density was decreased in Fmr1-cKO male mice. **a**, Representative images showing Iba1 staining in the IC of control and Fmr1-cKO mice at P14. Scale bar, 40 μm. **b**, Microglia size and density quantified in the IC of control and Fmr1-cKO mice at P14. Two-way ANOVA followed with Bonferroni’s multiple comparisons test, *p<0.05, **p<0.01, and ****p<0.0001, n=5 mice per group. **c**, qPCR quantification of mRNAs from inflammatory and homeostatic genes in microglia isolated from control and Fmr1-cKO mice at P12-14. Two-tailed unpaired t test, n=6 for control and 5 for Fmr1-cKO. **d**, Representative images showing PV staining in the IC of control and Fmr1-cKO at P22. Scale bar, 100 μm. **e**, Density of PV neurons in the IC was decreased in male but not in female Fmr1-cKO mice at P22. Two-way ANOVA followed with Bonferroni’s multiple comparisons test, **p<0.01, ns=not significant, n=5 mice per group. **f**, Density of PV neurons in the IC of control and Fmr1-cKO male mice at P14, P22 and P42. Two-way ANOVA followed with Bonferroni’s multiple comparisons test, ***p<0.001 and ****p<0.0001, n=5-6 mice per group. **g**, Total number of PV neurons in an IC was decreased in Fmr1-cKO male mice at P22 and P42 but not P14. Two-way ANOVA followed with Bonferroni’s multiple comparisons test, **p<0.01, n=4-5 mice per group. **h**, PV neuron density in the somatosensory cortex (SSC) was decreased in Fmr1-cKO males but not females. Two-way ANOVA followed with Bonferroni’s multiple comparisons test, ***p<0.001, ns = not significant, n=5 mice per group. **i**, PV neuron density in the IC of *Fmr1* KO male mice was decreased. Two-tailed unpaired t test, **p<0.01, n=5 mice per group. **j**, PV neuron density in the SSC of *Fmr1* KO male mice was decreased. Two-tailed unpaired t test, *p<0.05, n=3-4 mice per group.

To probe the increased density of microglia in the IC of male Fmr1-cKO mice, we examined Ki67-marked proliferating microglia at P10 (Extended Data Fig. 3e). There was a trend that a larger portion of microglia in the IC of male Fmr1-cKO mice were proliferating (Extended Data Fig. 3d). Furthermore, the reverse correction between the portion of proliferating microglia and the density of microglia was disrupted in the IC of male Fmr1-cKO mice (Extended Data Fig. 3f). This indicates that the increased microglia density in the IC of male mutant mice is a result of dysregulated proliferation.

In light of the microglial enlargement, we determined whether microglia became proinflammatory in male Fmr1-cKO mice. We purified microglia from the whole brain, because microglia in all brain regions we examined were larger in male Fmr1-cKO mice (Fig 2b and Extended Data Fig. 3b-c) and microglia from the IC were too limited. Quantitative RT-PCR analysis revealed normal expression of several proinflammatory genes in microglia of male Fmr1-cKO mice at P12-14: *C1qa* (encoding complement C1q subunit A), *Itgam* (encoding complement receptor 3), *Tnfa* (encoding tumor necrosis factor alpha), *Il1a* (encoding interleukin-1 alpha), and *Spp1* (secreted phosphoprotein 1) (Fig. 2c). These results suggest that microglia in Fmr1-cKO mice are not proinflammatory despite of their morphological changes.

### Reduced density of PV neurons in male Fmr1-cKO mice

As PV neurons exert powerful inhibition in the brain^24^, we reasoned that ablation of FMRP expression in microglia may result in a loss of PV neurons in the IC and thereby increase the susceptibility to AGS. We employed a stereological method to count immunoreactive cells on brain sections stained with antibodies against PV (Fig. 2d) and found reduced density of PV neurons in the IC of male, but not female, Fmr1-cKO mice at P22 (Fig. 2e). The reduction was not detected at P14 and was comparable at P22 and P42 (Fig. 2f), indicating that PV neurons are reduced during the 3^rd^ postnatal week. As the volume of the IC was the same in control and Fmr1-cKO mice at P14–P42 (Extended Data Fig. 3g), the total number of PV neurons in the IC was decreased to the same extent as the density in male Fmr1-cKO mice at P22 and P42 (Fig. 2g). Similarly, reduced density of PV neurons was observed in the somatosensory cortex (SSC) of male, but not female, Fmr1-cKO mice at P22 (Fig. 2h). In contrast, the density of PV neurons in the IC of male *Cx3cr1^CreER/+^* mice was not altered (Extended Data Fig. 3h), indicating that the mutation in one copy of the *Cx3cr1* gene in Fmr1-cKO mice does not contribute to the PV neuron phenotype. We observed reduced density of PV neurons in the SSC of male *Fmr1* null mice (Fig. 2j), as reported in previous studies^25,26^, and a similar phenotype in the IC (Fig. 2i). Taken together, these results show that FMRP deficiency in microglia is sufficient to cause reduced density of PV neurons in male mice. Because PV neuron density reduction only occurs in males, we used male mice in the remaining experiments.

### Single-nuclei RNA sequencing of the IC

To gain insight into the molecular mechanism on how FMRP deficiency in microglia caused reduced PV neuron density, we performed single-nuclei RNA sequencing (snRNA-seq) of IC tissues from male control and Fmr1-cKO mice at P22. Nuclei passing the quality check were used in analysis (Extended Data Fig. 4a-b). We performed principal component analysis and dimensional reduction using uniform manifold approximation and projection (UMAP) (Fig. 3a). By examining the expression of marker genes, we identified major cell types in the brain, including excitatory neurons, inhibitory neurons, astrocytes, oligodendrocytes, oligodendrocyte precursor cells (OPCs), and microglia (Fig. 3a-b and Extended Data Fig 4a-b). The differentially expressed genes were then subjected to Ingenuity Pathway Analysis (IPA). Interestingly, the pathways regulating lysosomal biogenesis were among the top enriched pathways in microglia, including CLEAR signaling, AMPK signaling and PPAR signaling (Fig. 3c-d). Differentially expressed genes in these pathways encode either important lysosomal enzymes (e.g., *Hexb, Ctsd,* and *Ctsf*)^61–63^ or signaling molecules essential for lysosome biogenesis (e.g., *Rptor, Ppp2r3a,* and *Tlr3*)^64,65^. This indicates that FMRP regulates lysosomal function, an understudied function for FMRP. Meanwhile, protein ubiquitination and BAG2 signaling pathways were enriched in Fmr1-cKO microglia (Fig. 3c, e). This suggests that the loss of FMRP impairs protein homeostasis in microglia, similar to that reported for neurons of *Fmr1^-/y^* mice^57^.

**Fig. 3:**
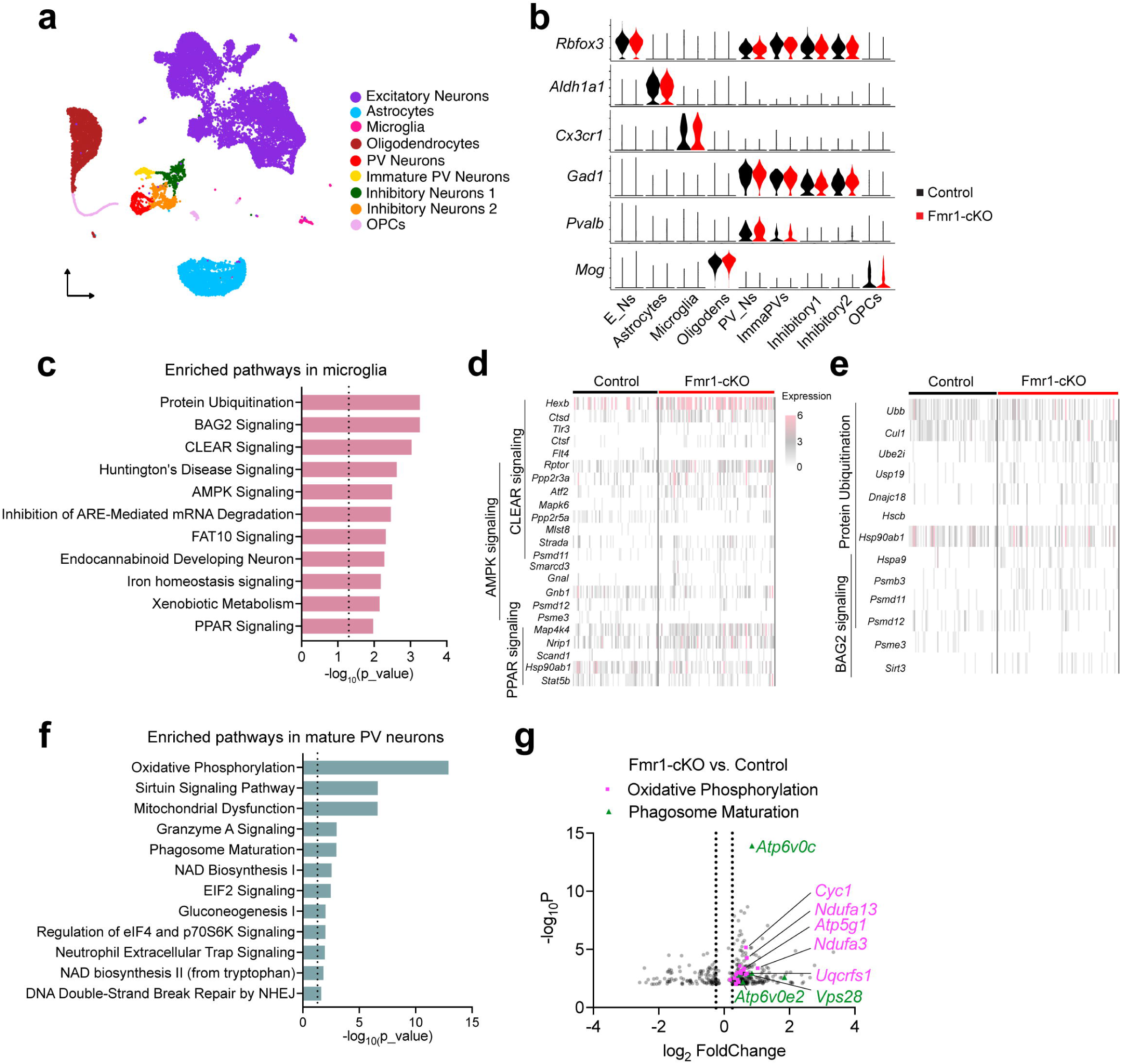
Single-nuclei RNA sequencing of the IC. **a**, UMAP plot of combined cells from the IC of male control and Fmr1-cKO mice at P22. Cell types were color-coded. **b**, Violin plots showing the expression of marker genes that were used to label different cell types. E_Ns, excitatory neurons; Oligodens, oligodendrocytes; PV_Ns, PV neurons; ImmaPVs: immature PV neurons. **c**, Top enriched signaling pathways in microglia of Fmr1-cKO mice. Dotted line indicates p=0.05. **d**, Heatmap showing differentially expressed genes associated with lysosomal pathways in microglia. **e**, Heatmap showing differentially expressed genes associated with protein degradation pathways in microglia. **f**, Top enriched signaling pathways in mature PV neurons of Fmr1-cKO mice. Dotted line indicated p=0.05. **g**, Volcano plot displaying differentially expressed genes in PV neurons. Genes belonging to the oxidative phosphorylation pathway are highlighted in magenta, while genes in the phagosome maturation pathway are showed in dark green. Dashed lines indicate log_2_FoldChange of ±0.25.

There are two clusters of PV neurons, mature PV neurons and immature PV neurons (Fig. 3a-b). In mature PV neurons of Fmr1-cKO mice, the top three enriched pathways were mitochondria-associated pathways, with the oxidative phosphorylation pathway exhibited the highest enrichment (Fig. 3f). Genes encoding members of the mitochondrial respiratory chain (e.g., *Cyc1, Ndufa13, Atp5g1,* and *Uqcrfs1*) were upregulated (Fig. 3g). Phagosome maturation was also among the top enriched pathways in Fmr1-cKO PV neurons at P22 (Fig. 3f and Extended Data Fig. 4c). In this pathway, upregulated genes included *Atp6v0c* and *Atp6v0e2*, which encode subunits of the vacuolar ATPase and are responsible for lysosomal acidification, and *Vps28*, which is involved in endo-lysosomal trafficking (Fig. 3g). These snRNA-Seq findings led us to ask how *Fmr1* deletion in microglia leads to transcriptomic changes in PV neurons and whether lysosomal and mitochondrial alterations contribute to the reduced density of PV neurons in Fmr1-cKO mice.

### FMRP regulates lysosomal gene expression in microglia

To unravel how FMRP deficiency leads to the transcriptomic alterations in microglia and PV neurons, we sought to identify FMRP binding targets in microglia. We immunoprecipitated whole brain lysates from male WT and *Fmr1*^-/y^ mice at P7-10 with specific FMRP antibodies (Fig. 4a), followed by quantitative RT-PCR, with a focus on lysosomal and microglial genes, because genes associated with the lysosome are among the most affected in Fmr1-cKO microglia. We measured FMRP binding to mRNAs encoding 5 lysosomal proteins (*Grn*, *Cst3*, *Ctsd*, *Hexb* and *Cd68*) (Fig. 4b) and several microglial proteins (Fig. 4c) with *Map1b* mRNA as the positive control. As shown previously^13^, FMRP immunoprecipitation pulled down *Map1b* mRNA from WT brain lysates (Fig. 4b). The immunoprecipitation also pulled down *Grn*, *Cst3*, *Ctsd*, *Hexb*, *Tyro3*, *Csf1r*, *Spp1*, *Aif1*, and *Atp8a2* mRNAs but not *Cd68*, *Trem2*, *C1qa*, *Tmem119*, *Mertk*, *Cx3cr1*, *Itgam*, and *Lgals9* mRNAs (Fig. 4b-c). While some of these targets were detected in previous high-throughput CLIP sequencing studies^66,67^ and *Grn* mRNA has been reported as a FMRP binding target through immunoprecipitation^68^, our data uncovered a distinct set of FMRP binding targets in microglia compared with other cell types.

**Fig. 4:**
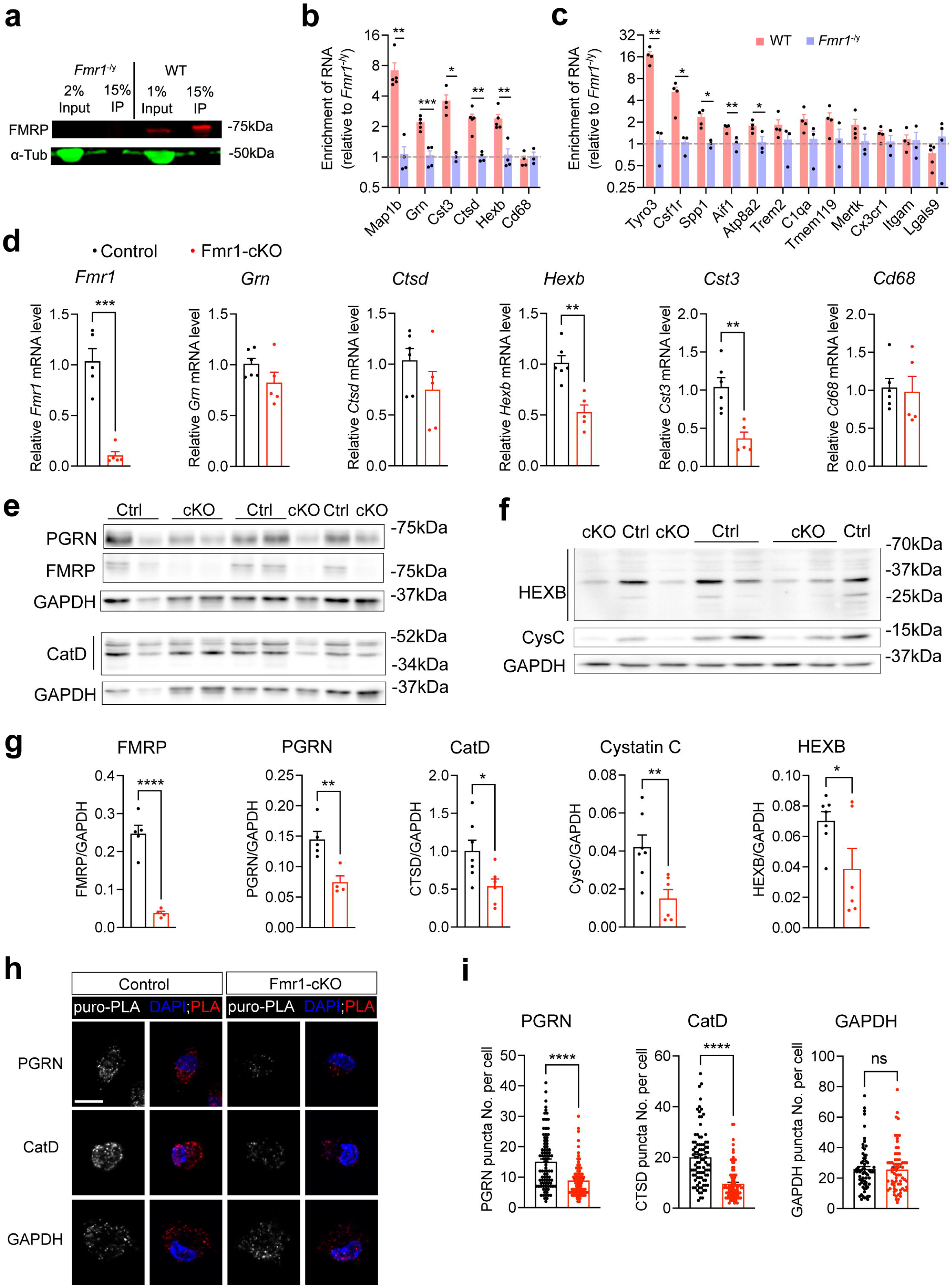
FMRP regulates expression of lysosomal proteins in microglia. **a**, Western blot showing that FMRP was specifically pulled down from the WT brain lysate by immunoprecipitation, while FMRP was absent in either *Fmr1^-/y^*brain lysate or immunoprecipitate. **b**, qPCR quantification of mRNAs encoding lysosomal proteins in FMRP immunoprecipitates. Two-tailed unpaired t test, *<0.05, **p<0.01, and ***p<0.001, n=3-5 male mice per group. **c**, qPCR quantification of microglial mRNAs in FMRP immunoprecipitates. Two-tailed unpaired t test, *<0.05 and **p<0.01, n=3-5 male mice per group. **d**, qPCR quantification of mRNAs encoding lysosomal proteins in microglia isolated from male control and Fmr1-cKO mice at P12-14. Two-tailed unpaired t test, **p<0.01, n=6 and 5 mice for control and Fmr1-cKO, respectively. **e** and **f,** Western blots showing protein levels of PGRN, FMRP, CatD (**e**), Cystatin C(CysC) and HEXB (**f**) in microglia isolated from male Control (Ctrl) and Fmr1-cKO (cKO) mice at P12-14. **g**, Quantification of protein levels in isolated microglia. Two-tailed unpaired t test, *p<0.05, **p<0.01, and ****p<0.0001, n=4-7 per group. **h**, Representative images showing PLA for puromycin with PGRN, CatD, or GAPDH in microglia isolated from male Control and Fmr1-cKO mice. Only proteins with incorporated puromycin will give PLA signals. Scale bar, 10 μm. **i**, Quantification of PLA puncta number for puromycin with PGRN, CatD, or GAPDH in each microglia. Two-tailed unpaired t test, ****p<0.0001, ns=not significant, n=30 cell per mouse, 3 mice per genotype.

We next sought to explore how *Fmr1* deletion alters the expression of FMRP binding targets in microglia. We purified microglia from male Fmr1-cKO and control mice at P14 and quantified levels of these mRNA targets and their proteins. While levels of *Spp1*, *Grn* and *Ctsd* mRNAs were not altered (Fig. 2c and 4d), levels of *Hexb and Cst3* mRNAs were reduced in Fmr1-cKO microglia (Fig. 4d). Immunoblotting analysis showed approximately 50% reductions in the protein levels of the four lysosomal targets: progranulin (PGRN, encoded by the *Grn* gene), cathepsin D (CatD, encoded by the *Ctsd* gene), Cystatin C (CysC, encoded by the *Cst3* gene) and hexosaminidase B (HEXB, encoded by the *Hexb* gene) in Fmr1-cKO microglia (Fig. 4e-g). As a validation of gene deletion, *Fmr1* mRNA and FMRP were hardly detectable in Fmr1-cKO microglia (Fig. 4d, e and g).

Since levels of *Grn* and *Ctsd* mRNAs were not changed but levels of their translation products were decreased by ∼50% in Fmr1-cKO microglia, we hypothesized that FMRP enhances translation of *Grn* and *Ctsd* mRNAs. To test this hypothesis, we adopted puromycin-mediated proximity ligation assay (PLA) to quantify the translation rate in cultured microglia isolated from male control and Fmr1-cKO mice at P12-14 (Extended Data Fig. 5a). Puromycin labeled newly synthesized polypeptides, and a PLA signal was produced when the antibody against puromycin and the antibody against PGRN, CatD, or GAPDH bound to the same polypeptides. Synthesis rates of PGRN and CatD were both decreased in microglia of Fmr1-cKO mice, as reflected by the decreased number of PLA puncta, while the synthesis rate of GAPDH was not affected as *Gapdh* mRNA is not a target of FMRP (Fig. 4h-i). Taken together, these results indicate that FMRP facilitates the expression of lysosomal proteins by increasing the abundance (*Hexb* and *Cst3*) and translation (*Grn* and *Ctsd*) of mRNAs in microglia.

### Lysosomal function was impaired in microglia with *Fmr1* deficiency

We reasoned that reduced expression of lysosomal proteins due to FMRP deficiency should impair lysosomal function in microglia. We first examined microglial lysosomal morphology in the IC of male Fmr1-cKO mice, as lysosomes become enlarged due to the buildup of undigested substances when they are dysfunctional^69^. Immunostaining of microglial lysosomal marker CD68 revealed significant increases in lysosomal volume and number per microglia in Fmr1-cKO mice (Fig. 5a-b). Lysosomal number was increased across all lysosomal sizes, especially the largest-size group (Fig. 5c). Consistently, microglia in the IC of male *Fmr1* null mice exhibited similar increases in lysosomal volume and number (Fig. 5d-f).

**Fig. 5:**
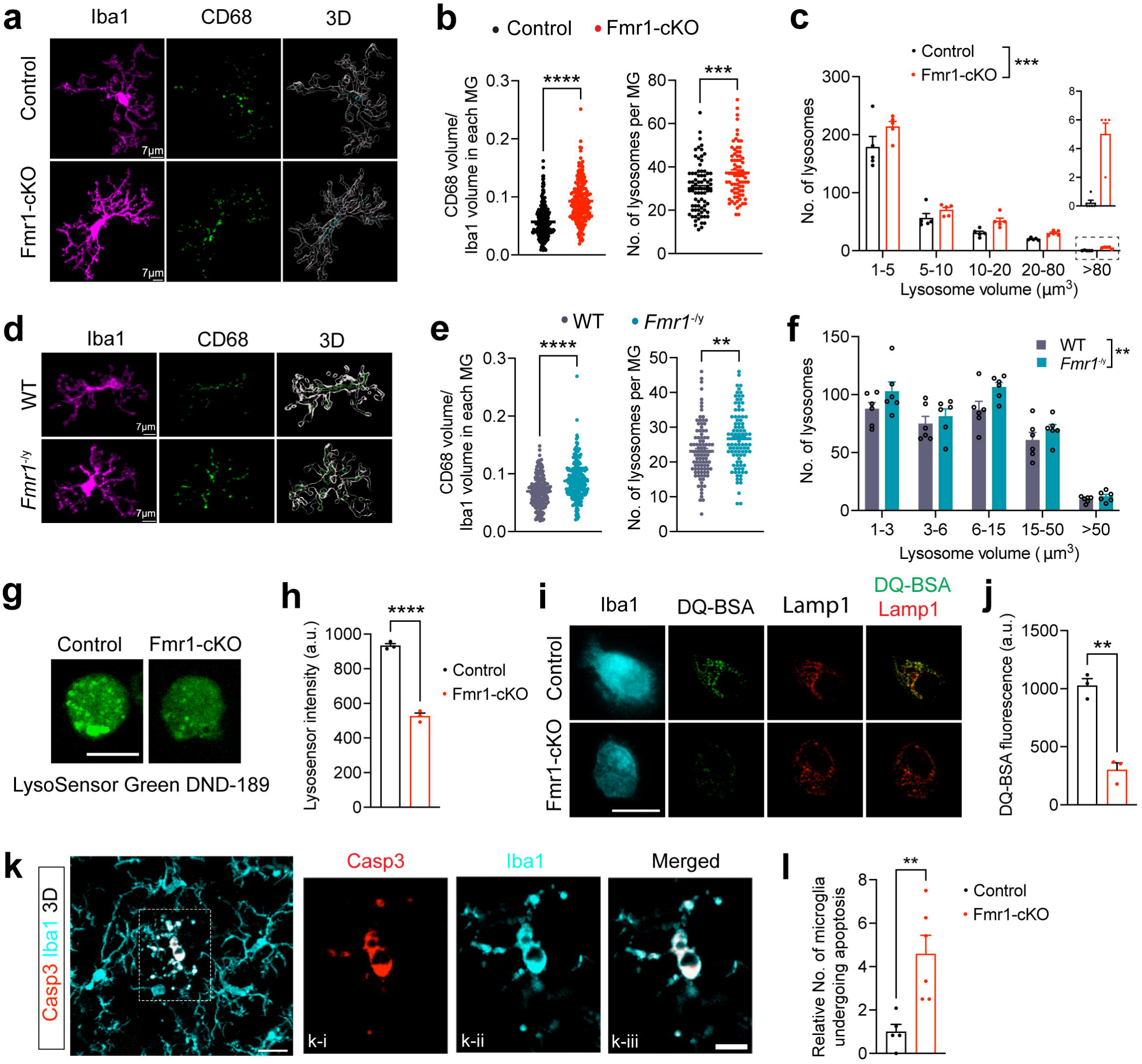
Lysosomal function was impaired in microglia with *Fmr1* deficiency. **a**, Fluorescent immunohistochemistry of Iba1 and CD68 in the IC of male control and Fmr1-cKO mice at P14. Z stack of confocal images are used to generate 3D models of microglia and lysosomes with Imaris software. **b**, Normalized CD68 volume and the number of CD68 puncta per microglia were increased in IC of Fmr1-cKO mice. Two-tailed unpaired t test. ***p<0.001 and ****p<0.0001, n=5 male mice per group. Approximately 50 microglia per mouse were measured for CD68 volume, and 13-16 microglia per mouse were measured for lysosome numbers. **c**, Distribution of lysosomes with different volumes. Two-way ANOVA followed with Bonferroni’s multiple comparisons test, ***p<0.001, n=5 male mice per group. Lysosomes in 14-16 microglia per mouse were analyzed. **d**, Fluorescent IHC of Iba1 and CD68 in the IC of WT and *Fmr1^-/y^* mice at P14. Confocal z stack images were used to generate 3D models with Imaris software. **e**, Relative volume and the number of CD68^+^ puncta per microglia were increased in the IC of *Fmr1^-/y^* mice at P14. Two-tailed unpaired t test. Left graph: ****p<0.0001, n=5 mice per group and ∼50 microglia per mouse. Right graph: **p<0.01, n=5 mice per group and 15 microglia in IC per mouse. **f**, Distribution of lysosomes with different volumes. Two-way ANOVA followed with Bonferroni’s multiple comparisons test, **p<0.01, n=5 male mice per genotype and 16 microglia per mouse. **g**, Representative confocal images showing LysoSensor Green DND-189 fluorescence in microglia isolated from control and Fmr1 cKO mice at P12-14. Scale bar, 10 μm. **h**, Intensity of LysoSensor fluorescence was significantly decreased in Fmr1-cKO microglia. Two-tailed unpaired t test, ****p<0.0001, n=3 male mice per genotype and 40 microglia per mouse. **i**, Super-resolution microscopic images showing that DQ-BSA fluorescence (green) were colocalized with Lamp1+(red) lysosomes in Iba1+(cyan) microglia, isolated from Control and Fmr1-cKO mice at P12-14. Scale bar, 10 μm. **j**, DQ-BSA fluorescence was decreased in Fmr1-cKO microglia. Two-tailed unpaired t test, **p<0.01, n=3 male mice per genotype and 20-30 microglia per mouse. **k**, 3D image showing apoptotic microglia in the IC of male Fmr1-cKO mice at P14. The outlined area is enlarged in single focal plane images to show a microglia positive for activated Casp3 in k-i, k-ii and k-iii. scale bars, 15 μm in k and 8 μm in k-iii. **l**, Relative number of apoptotic microglia were increased in the IC of male Fmr1-cKO mice. Two-tailed unpaired t test, **p<0.01, n=5 for control and 6 for Fmr1-cKO.

The lysosome is an acidic organelle, and the high acidity is required for activation of lysosomal hydrolases^69^. To quantify the acidity of lysosomes in microglia, we used LysoSensor Green DND-189, a pH-sensitive dye that can be absorbed by acidic organelles and becomes more fluorescent in acidic environments. Microglia isolated from Fmr1-cKO mice showed a drastic decrease in intensity of LysoSensor fluorescence (Fig. 5g-h). To assess the digestive capability of lysosomes, we utilized the substrate DQ-BSA, which can be taken up by microglia and is converted to brightly fluorescent products upon hydrolysis in lysosomes. Super-resolution microscopy confirmed that DQ-BSA puncta were colocalized with lysosomal marker Lamp1 in Iba1^+^ microglia (Fig. 5i and Extended Data Fig. 5b). Microglia isolated from Fmr1-cKO mice exhibited decreased capacity of breaking down DQ-BSA compared with control microglia (Fig. 5i-j). Similarly, lysosomal acidity and degradation capability were both decreased in microglia purified from *Fmr1* null mice at P12-14 compared with microglia from WT littermates (Extended Data Fig. 5c-f). Collectively, these results indicate that FMRP deficiency impairs the ability of lysosomes to break down substances in microglia.

Accumulation of undegraded substrates within lysosomes leads to cell death in lysosomal storage diseases^70^. Our immunohistochemistry with antibodies against Iba1 and cleaved caspase 3, a marker for apoptosis, on brain sections revealed a drastic increase in apoptotic microglia in male Fmr1-cKO mice at P14 compared with control littermates (Fig.5k-l). This suggests that the impairment in lysosomal function due to FMRP deficiency causes death of microglia through apoptosis.

### PV neurons undergo apoptosis in response to progranulin insufficiency

Homozygous loss-of-function (LOF) *GRN* mutations result in a neuronal ceroid lipofuscinosis, while heterozygous LOF *GRN* mutations lead to frontotemporal lobe degeneration (FTLD-GRN)^71^. Given that PGRN is largely expressed in microglia^52^ (Extended Data Fig. 6a) and regulates acidification and trafficking of lysosomes^48,49^, we hypothesized that PGRN insufficiency results in lysosomal dysfunction and increased apoptosis in PV neurons. As impaired lysosomal function is known to lead to mitochondrial dysfunction in neurons^72^, this hypothesis is further supported by our snRNA-Seq data, which shows mitochondrial dysfunction in PV neurons of male Fmr1-cKO mice (Fig. 3f). PV neurons express two PGRN receptors^52^, *Sort1* and *Lrp1* (Extended Data Fig. 6a) and thus are able to internalize extracellular PGRN to support their lysosomal function and survival.

To test our hypothesis, we first quantified the number of apoptotic PV neurons by carrying out florescent immunohistochemistry with antibodies against PV and cleaved caspase 3 in the IC of male control and Fmr1-cKO mice at P14. The number of apoptotic PV neurons was increased by ∼6 fold in Fmr1-cKO mice compared with control mice (Fig. 6a-b). We then examined lysosomes in PV neurons in the IC of male mice at P14 using the lysosome marker Lamp1 (Fig. 6c). Lamp1-marked lysosomes were enlarged in PV neurons in the IC of Fmr1-cKO mice compared with those in control mice (Fig. 6c-d and Extended Data Fig. 6b-d), indicative of lysosomal dysfunction. One hallmark in FTLD-GRN is accumulation of phosphorylated transactivation response DNA-binding protein 43 (pTDP-43) in the cytoplasm^73^. We found that there were more numerous large pTDP-43 aggregates in the IC in Fmr1-cKO mice than in control mice at P14 (Fig. 6e-g). The number of apoptotic cells associated with pTDP-43 aggregates was increased by ∼20 fold in Fmr1-cKO mice (Fig. 6h), including apoptotic microglia (Fig. 6e) and apoptotic PV neurons (Fig. 6f). These results strongly suggest that PGRN insufficiency is the main cause for reduced density of PV neurons in male Fmr1-cKO mice.

**Fig. 6:**
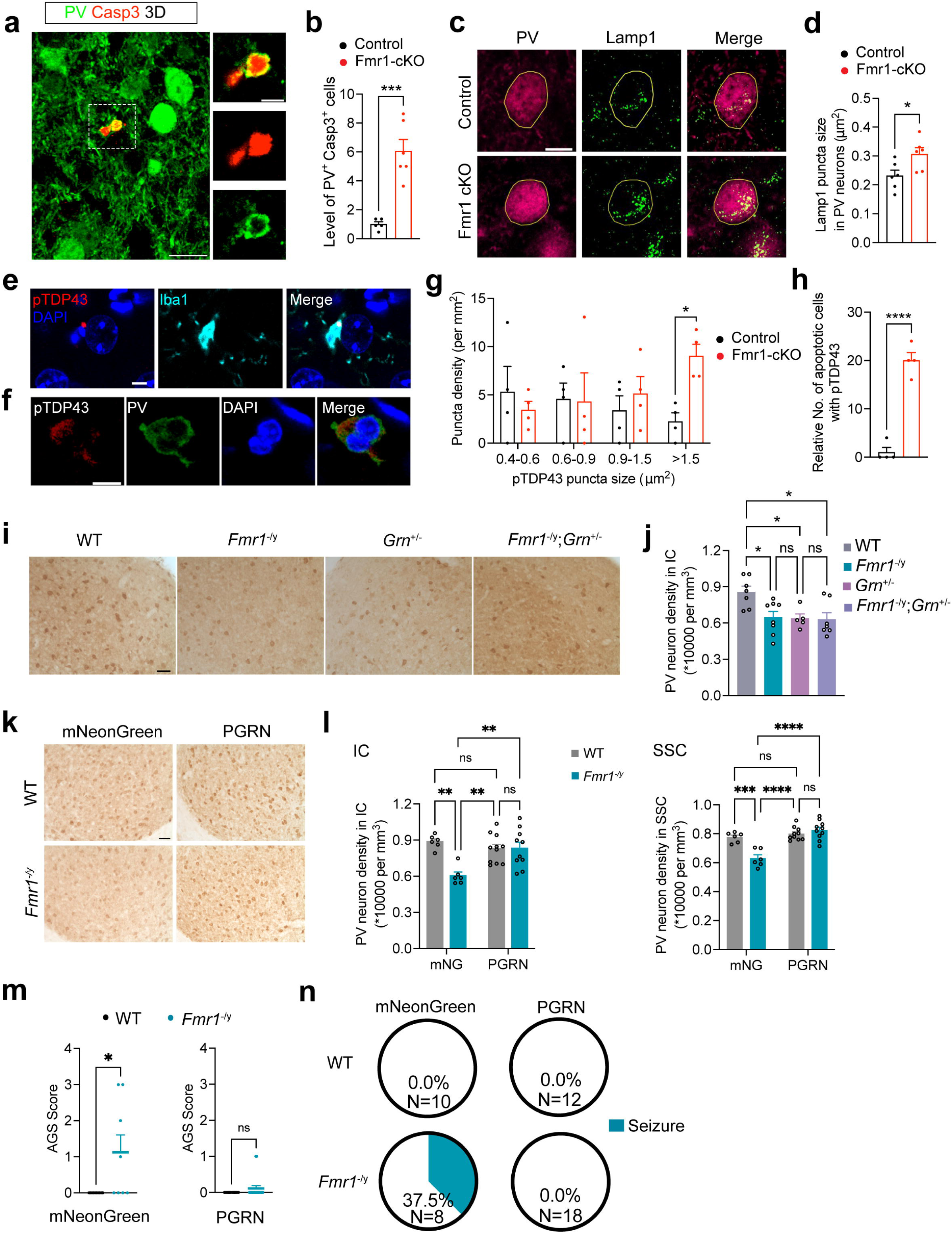
PV neurons undergo apoptosis in response to progranulin insufficiency. **a**, 3D image reconstructed from confocal z stack serials showing a PV neuron undergoing apoptosis in the IC of Fmr1-cKO male mice at P14. Smaller images on the right side are enlarged from the outlined area and are single focal plane images showing the colocalization of PV and cleaved caspase 3 (Casp3). Scale bars, 20 μm in the 3D image and 5 μm in the side images. **b**, Relative number of apoptotic PV neurons was significantly increased in the IC of male Fmr1-cKO mice. Two-tailed unpaired t test, ***p<0.001, n=5 for Control and 6 for Fmr1-cKO. **c**, Immunohistochemistry of PV and Lamp1 in the IC of male Control and Fmr1-cKO mice at P14. Golden lines outline the soma of a single PV neuron. Scale bar, 10 μm. **d**, Size of Lamp1^+^ puncta in PV neurons was significantly increased in male Fmr1-cKO mice. Two-tailed unpaired t test, *p<0.05, n=6 mice per genotype and 10 PV neurons per mouse. **e**, Confocal images showing immunoreactivity of pTDP43 in an apoptotic microglia of male Fmr1-cKO mice at P14. Scale bars, 5 μm. **f**, Confocal images showing immunoreactivity of pTDP43 in an apoptotic PV neuron of male Fmr1-cKO mice at P14. Scale bars, 5 μm. **g**, Density of pTDP43 puncta across a range of different sizes in the IC of male mice at P14. Two-way ANOVA followed with Bonferroni’s multiple comparisons test, *p<0.05, n=4 mice per genotype. **h**, Relative number of apoptotic cells positive for pTDP43 puncta was increased in male Fmr1-cKO mice. Two-tailed unpaired t test, ****p<0.0001, n=4 mice per genotype. **i**, Representative images of PV immunohistochemistry in the IC of male WT, *Fmr1^-/y^*, *Grn^+/-^*, and *Fmr1^-/y^*;*Grn^+/-^*mice at P22. Scale bar, 100 μm. **j**, PV neuron density in the IC of male WT, *Fmr1^-/y^*, *Grn^+/-^*, and *Fmr1^-/y^*;*Grn^+/-^*mice at P22. One-way ANOVA followed with Tukey’s multiple comparisons test, ns, not significant and *p<0.05, n=5-8 mice per group. **k**, Representative images of PV immunohistochemistry in the IC of P22 male WT and *Fmr1*^-/y^ mice overexpressing mNeonGreen or progranulin (PGRN). Scale bar, 50 μm. **l**, Density of PV neurons in the IC and SSC of male WT and *Fmr1*^-/y^ mice overexpressing mNeonGreen (mNG) or PGRN. Two-way ANOVA followed with Tukey’s multiple comparisons test, **p<0.01, ***p<0.001, ****p<0.0001, and ns = not significant. n=10-11 for PGRN-overexpressing WT and *Fmr1*^-/y^ mice, n=6 for mNeonGreen-overexpressing WT and *Fmr1*^-/y^ mice. **m**, AGS score of male WT and *Fmr1*^-/y^ mice overexpressing mNeonGreen or PGRN. Mann-Whitney test, *p<0.05, ns = not significant. Sample sizes are shown in panel (**n**). **n**, Pie graphs showing the percentage of mice developed seizure (AGS score >=2) in each group.

We next asked if progranulin insufficiency alone could reduce the density of PV neurons. To this end, we crossed female *Fmr1^+/-^* mice with male *Grn^+/-^* mice to produce male mice of four genotypes: WT, *Fmr1^-/y^*, *Grn^+/-^*, and *Fmr1^-/y^;Grn^+/-^*. We measured the density of PV neurons in the IC of these mice at P22. The density of PV neurons was reduced in *Grn^+/-^*, *Fmr1^-/y^*, and *Fmr1^-/y^;Grn^+/-^* mice to a similar extent (Fig. 6i-j), suggesting that FMRP deficiency and progranulin insufficiency cause apoptosis of PV neurons through the same pathway. Taken together, these data show that progranulin downregulation in microglia is the major contributor to the reduced density of PV neurons in *Fmr1* null mice. This prompted us to ask if increasing progranulin level in the brain of *Fmr1* null mice is able to reverse the reduction in PV neurons.

To answer this question, we generated AAV8-CAG-Grn for progranulin expression and control AAV8-CAG-mNG for mNeonGreen expression (Extended Data Fig. 7a-c). Following intracerebroventricular injection into the lateral ventricle at P0, the AAV8 viruses spread throughout the whole brain (Extended Data Fig. 7e). Viral progranulin overexpression in the IC and SSC was validated by western blot (Extended Data Fig. 7d) and immunohistochemistry (Extended Data Fig. 7f). We then injected AAV8 viruses expressing either mNeonGreen or progranulin into the lateral ventricle of male *Fmr1* null mice and WT littermates at P0. While mNeonGreen overexpression did not affect reduced density of PV neurons in the IC and SSC of *Fmr1* null mice, progranulin overexpression completely normalized the density of PV neurons (Fig. 6k-l). In addition, progranulin overexpression prevented *Fmr1* null mice from AGS, with 0 out of 18 mice developing seizures, compared to 37.5% of mNeonGreen-expressing *Fmr1* null mice (3 out of 8 mice) developing seizures (Fig. 6m-n). These results suggest that elevating progranulin is a potent therapeutic intervention for FXS.

## Discussion

Reduced number of PV neurons has been consistently observed in multiple cortical regions of FXS patients and in the cortex of the FXS mouse model (*Fmr1* null mice)^24–27^. This study also found reduced number of PV neurons in the IC of *Fmr1* null mice, suggesting that this PV neuron abnormality is not restricted to the cortex of FXS patients. By selectively ablating FMRP expression in microglia postnatally, we found that this conditional mouse knockout, Fmr1-cKO, recapitulated this PV neuron abnormality as well as AGS, another phenotype that is consistently observed in *Fmr1* null mice^57,58^. Given that individual PV neurons can innervate more than 1,000 excitatory pyramidal neurons^36^, the loss of PV neurons in the IC or other parts of the auditory pathway is the likely cause for AGS in *Fmr1* null mice and Fmr1-cKO mice as well as seizures in FXS patients. Reduced number or hypoactivity of PV neurons has also been observed in postmortem tissues from individuals with idiopathic ASD^28,29^ and other ASD mouse models^30–35^. It is important to determine whether microglial dysfunction is also the cause for the PV neuron abnormality in these cases in future studies.

Previous studies using high-throughput CLIP-seq have identified hundreds of FMRP targets, a high percentage of which encodes axonal and dendritic proteins^11,66,74^. However, these studies used whole brain tissues, which makes it less successful to identify the targets expressed in low abundance of cell types such as microglia. Based on the snRNA-Seq data from Fmr1-cKO mice, we employed a candidate approach with a focus on mRNAs for lysosomal proteins. This approach allowed us to identify *Grn*, *Cst3*, *Ctsd* and *Hexb* mRNAs as FMRP targets. In contrast to its role in neurons where FMRP generally suppresses expression of target mRNAs^12–15,75^, FMRP appears to promote expression of its mRNA targets in microglia. For example, we found that FMRP stimulates translation of *Grn* and *Ctsd* mRNAs while elevating levels of *Cst3* and *Hexb* mRNAs. In the absence of FMRP, reduced expression of these genes would induce compensatory responses in transcription of the genes for these proteins and other lysosomal proteins. This could be the reason why several pathways related to lysosome biogenesis are enriched in the microglial snRNA-Seq dataset from Fmr1-cKO mice. Although the current study was focused on apoptosis of microglia and PV neurons, lysosomal impairments would also reduce the capacity of microglial phagocytosis. Given that microglia prune excessive synapses through phagocytosis in the postnatal brain development, further studies are needed to investigate whether and how FMRP loss affects phagocytic functions of microglia and remodeling of neural circuits.

Our study indicates that PGRN insufficiency is the main cause for reduced density of PV neurons in FXS. First, *Fmr1* deletion in microglia reduced the density of PV neurons in the IC and SSC to the same extent as observed in *Fmr1* null mice. Second, FMRP deficiency halved the expression of progranulin in microglia, which are the major source of progranulin in the brain. Third, PV neurons in Fmr1-cKO mice displayed enlarged lysosomes and pTDP-43 accumulation, two pathological features of progranulin haploinsufficiency^73^. Fourth, the density of PV neurons in the IC of *Fmr1^-/y^* null mice was reduced to the same extent as in the IC of male *Grn^+/-^* mice or *Fmr1^-/y^;Grn^+/-^*mice. Finally, viral overexpression of progranulin prevented the loss of PV neurons in the IC and SSC of *Fmr1^-/y^*null mice. One exception to this otherwise impeccable evidence is that the density of PV neurons in the SSC of *Grn^+/-^*mice was not significantly reduced (Extended Data Fig. 6e). This suggests that there are additional unknown factors in FMRP-deficient microglia that contribute to death of PV neurons in FXS, and thus progranulin insufficiency is not the only cause for the loss of PV neurons. Our study also provides an explanation why the neuronal loss in the cortex of FXS patients is largely specific to PV neurons^27^. Damaged mitochondria are removed through the autophagy-lysosome pathway^76^. As PV neurons have high demands on mitochondria due to the fast-spiking properties^77^, their mitochondria are particularly susceptible to damages. If in FXS patients and mouse models lysosomes in PV neurons are dysfunctional due to progranulin insufficiency, damaged mitochondria would not be removed efficiently and are accumulated, leading to apoptosis of PV neurons. This could be the reason why mitochondrial dysfunction is associated with neurodegeneration in lysosomal storage diseases^72^.

Our study further shows that progranulin overexpression in the brain rescues AGS and the loss of PV neurons in *Fmr1* null mice. Importantly, several progranulin-elevating drugs are currently under investigation in clinical trials for treatment of FTLD-GRN^71^, and they could be repurposed for treatment of FXS patients and other ASD patients associated with reduced number or hypoactivity of PV neurons.

### Methods Mouse strains

The *Fmr1^lox^* mouse strain^78^ was obtained from Dr. David Nelson at Baylor College of Medicine. *Cx3cr1^CreER/+^* (Strain #: 020940), Ai9 (Strain #: 007909), *Grn*^-/-^ (Strain #: 013175), *Fmr1* KO (Strain #: 003025), and C57BL/6J (Strain #: 000664) mouse strains were obtained from the Jackson Laboratory. All mouse strains were maintained on the C57BL/6J genetic background. *Fmr1^lox/lox^* females were crossed with *Cx3cr1^CreER/+^* mice to generate male *Fmr1^lox/y^*;*Cx3cr1^CreER/+^*mice. Then, *Fmr1^lox/lox^* females were crossed to *Fmr1^lox/y^*;*Cx3cr1^CreER/+^* males, and their offspring were treated with 9 μl tamoxifen (Sigma, T5648, 40 mg/ml in corn oil) at P0 to generate control and Fmr1-cKO mice. We crossed female *Fmr1^+/-^* mice to male C57BL/6J WT mice to get male *Fmr1^-/y^* and WT mice. Mice were maintained in a standard mouse holding room with 12-h light-dark cycle and had free access to food and water. All animal procedures were approved by the UF Scripps IACUC (protocol #16-003) and conducted according to the guidelines.

### Immunohistochemistry

Pregnant females with embryos at E14.5 and E16.5 were euthanized by carbon dioxide asphyxiation. Embryonic brains and P0 brains were dissected out and fixed in 4% paraformaldehyde (PFA) for 7-8 h. For postnatal brains older than P0, mice were transcardially perfused with ice-cold PBS followed with 4% PFA in PBS. Brains were dissected and postfixed at 4°C in 4% PFA for 6-24 h, depending on the primary antibodies to be used. After fixation, the brains were transferred to 30% sucrose in PBS at 4°C for over 2 days until the brains sank to the bottom of the buffer. Embryonic brains and P0 brains were sectioned at the thickness of 20 μm with a Leica Cryostat CM3050. Postnatal brains were sectioned at thickness of 40 μm with a Leica microtome SM2010R. Sections were stored at -20°C in 3:3:3:1 ethyleneglycol / glycerol / H_2_O / 10XPBS until immunohistochemistry.

For fluorescent immunohistochemistry, sections were washed in PBS and then blocked in PBS with 10% normal horse serum and 0.3% Triton X-100 for 2 h. Sections were then incubated at 4°C overnight with primary antibodies diluted in PBS with 1% normal horse serum and 0.3% Trion X-100. Primary antibodies against Iba1 (Wako, 019-19741, 1:500), Iba1 (Synaptic System, 234 308, 1:500), FMRP (abcam, ab17722, 1:300), parvalbumin (Millipore Sigma, P3088, 1:2000), Ki67 (abcam, ab15580, 1:500), cleaved caspase-3 (Cell Signaling, #9661, 1;400), CD68 (Bio-rad, MCA1957, 1:400), Lamp1 (DSHB, 1D4B, 1:250), and pTDP43 (Proteintech, 80007-1-RR, 1:400) were used. Secondary antibodies were diluted in the same diluent as primary antibodies and incubated at room temperature for 1.5 h. Secondary antibodies were from Jackson ImmunoResarch: Alexa Fluro 488 donkey anti-guinea pig (#706-545-148), Alexa Fluro 594 donkey anti-guinea pig (#706-585-148), Alexa Fluro 647 donkey anti-guinea pig (#706-605-148), Alexa Fluro 594 donkey anti-rabbit (#711-585-152), Alexa Fluro 488 donkey anti-rabbit (#711-545-152), Alexa Fluro 488 donkey anti-mouse (#715-005-150), Alexa Fluro 594 donkey anti-mouse (#715-605-150), and Alexa Fluro 488 Donkey anti-rat (#712-545-150). Sections were incubated with DAPI (2.5 μg/ml) in PBS for 5 min. Sections were mounted onto Superfrost plus slides, and Fluoro Gel mounting medium (Electron microscope science, 17985-10) was applied.

For 3,3′-diaminobenzidine (DAB) immunostaining, sections were washed with TBS and quenched with 10% methanol and 3% H_2_O_2_ in TBS for 18 min. After washing with TBS, the sections were blocked in TBS with 0.4% Triton X-100, 1% glycine, 1% BSA and 10% normal horse serum for 2 h. Primary antibodies against Iba1 (Wako, 019-19741, 1:500) or parvalbumin (Millipore Sigma, P3088, 1:2000) were diluted in blocking buffer and incubated with section at 4°C overnight. Sections were incubated with biotinylated secondary goat anti-rabbit (Vector, BA-1000, 1:200) or goat anti-mouse (Vector, BP-9200-50, 1:200) antibodies for 1 h at room temperature. The VECTASTAIN Elite ABC Kit (cat# PK-6100) were used for ABC reactions following the manufactory’s protocols. Briefly, buffer A and B were diluted at 1:100 in blocking buffer and mixed for 25 min at room temperature, followed by incubation with sections for 30 min at room temperature. After wash with TBS and 100 mM Tris-HCl (pH7.5), sections were developed with 0.05% DAB, 0.003% H_2_O_2_ in 100 mM Tris-HCl (pH7.5) for 3-5 min. After transferring onto Superfrost plus slides, sections were dehydrated by sequential incubation in 70% ethanol (5 min), 95% ethanol (5 min), 100% ethanol (twice, 5 min each), and xylene (twice, 10 min each). DPX Mountant and coverslip were applied.

### Audiogenic seizure test

Mice of both sexes at age of P22-25 were used for the test. The test was performed following a published protocol^79^ with minor modifications. Mice were handled for 3 consecutive days before the test was conducted. On the experimental day, mice were transferred to a quiet room (<60dB) for 30-min habituation and then to a testing chamber. After 1 min of habituation, a 128-dB white noise was applied for two 2-min trials separated by a 1-min inter-trial interval. Seizure behavior was scored as follows: 0, no response; 1, wild running; 2, clonic seizure; 3, tonic seizure; 4, respiratory arrest/death.

### Density and size of microglia

Every other sections of the IC from P14 mice were used for quantification of microglial density and size. Every third sections of the hippocampus and striatum from P14 mice were used for quantification of microglial density and size. DAB-stained sections were examined under a Nikon Eclipse E800 microscope. Stereo Investigator software (MBF Bioscience) was used to quantify density and size of Iba1-immunoreactive microglia in different brain regions. Both males and females were used in this experiment.

### Percentage of proliferating microglia

Every other sections of the IC from P10 male mice were used for fluorescent immunohistochemistry of Ki67 and Iba1. Ten images were randomly taken for each mouse under a Nikon C2 confocal microscope with a 20X lens. Microglia number per image was counted with the Nikon NIS-Element analyzer, and the cell density was calculated. Iba1 and Ki67 double positive cells were counted as proliferating microglia and the percentage of proliferating microglia were calculated for each image.

### PV neuron density

Every other sections of the IC from P14 mice and every third sections of the IC from P22 and P42 mice were used for quantification of PV neuron density. Every fourth sections of the somatosensory cortex from P22 mice were used to quantify PV neuron density. DAB immunostaining against parvalbumin was performed. Stereo Investigator software (MBF Bioscience) was used to quantify cell density. To calculate the IC volume, the area of the IC on each section was measured with Stereo Investigator. Then the IC volume for one hemisphere was estimated based on the number of sections, thickness of section and averaged area of IC per section. Total number of PV neurons was calculated based on cell density and IC volume.

### Microglia purification

Male mice at P7-14 were decapitated. Brains were dissected out and minced into small chunks (∼1 mm^3^). Brain tissues were dissociated with 20U/ml papain (Worthington, LS003126) in EBSS containing 0.7 mg/ml DNase I (Sigma, DN25) for 40-60 min on a shaker in a 37°C incubator. Brain tissues were triturated to release single cells by pipetting with a 5 ml serological pipette for a few rounds until no visible brain tissues existed. After passing through a 40-μm cell strainer, cells were pelleted by centrifugation at 300g for 8 min. EBSS containing ovomucoid protease inhibitor (Worthington, LK003182) was added to the pellet to resuspend cells. After centrifugation to pellet cells, a step for myelin removal was included for P12-14 mice^80^. Cells were resuspended in 6.7 ml DPBS (PBS containing 0.1 mg/ml DNase I). Cell suspension was then mixed with 2 ml myelin separation buffer: 1.8 ml Percoll (Sigma, P4937), 200 μl 10X DPBS, 1.8 μl 1M CaCl_2_, and 1 μl 1M MgCl_2_. Cells were collected by centrifugation at 500g for 15 min with slow brake. Microglia were purified using CD11b microbeads (Miltenyi Biotec, 130-093-634) according to manufacturer’s protocols. Briefly, cells were resuspended in 180 μl buffer (PBS with 0.5% BSA) and incubated with 20 μl CD11b microbeads at 4°C for 15 min. Cells were then washed and resuspended in the buffer. A LS column (Miltenyi Biotec, 130-042-401) was inserted in the magnetic field of a MidiMACS Separator (Miltenyi Biotec, 130-042-302). Cell suspensions were passed through the column. Microglia stayed in the column in the magnetic field, while the other types of cells flowed through the column. The column was taken out from the magnetic field, and microglia were collected by washing the column with buffer, followed by centrifugation.

### qPCR quantification of gene expression in microglia

Total RNAs of microglia were extracted with Absolutely RNA Nanoprep Kit (Agilent, 400753) according to the manufacturer’s protocol. Reverse transcription was done with SuperScript IV First-Strand synthesis system (Invitrogen, 18091050). qPCR was done with SYBR Green master mix (Applied Biosystems, A25742).

### Single nucleus preparation for snRNA sequencing

Male mice at P22 were decapitated. IC tissues were quickly dissected out, snap frozen in liquid nitrogen and stored at -80°C. IC tissues from 3 mice were combined for each genotype. The suspension of single nuclei was prepared following a published method^81^ with minor revisions. Frozen IC tissues were homogenized (Sigma D8938-1SET) in 2 ml EZ lysis buffer (Nuclei isolation kit: Sigma, NUC101) by 15 strokes of pestle A and 20 strokes of pestle B. The homogenate was topped to 4 ml with EZ lysis buffer and put on ice for 5 min. Lysates were centrifuged at 500g at 4°C for 5min, and the pellet was resuspended with 4 ml EZ lysis buffer. After 5 min incubation on ice, nuclei were centrifuged at 500g at 4°C for 5 min. The pellet of nuclei were resuspended in 3 ml of PBS suspension buffer: PBS with 1% BSA, 0.2 U/μl RNasin (Promega, N2611) and 0.2 U/μl SUPERase-IN RNase inhibitor (Invitrogen, AM2696), and were filtered through a 40-μm cell strainer. 45 μl of propidium iodide (BD DNA QC particles, 50 μg/ml) was added into 2 ml of nuclei suspension to label nuclei for sorting. Single nuclei were sorted on a BD FACSAria lII cell sorter, with a 100 μm nozzle. Sorted single nuclei were collected in a 1.5-ml EP tube with 200 μl of PBS suspension buffer with 4 μl of additional SUPERase-IN RNase inhibitor. The concentration of sorted nuclei was accessed and adjusted to 1000 nuclei per μl.

### snRNA sequencing

Single-cell libraries were generated using 10x Genomics Chromium Next GEM Single Cell 3’ Kit v3.1 reagents (P/N 1000269), according to the manufacturer’s recommendations. Briefly, for each sample a total of 16,500 nuclei were loaded on one lane of a Chip G Single Cell Kit (P/N 1000127). Single-cell gel beads in emulsion (GEMs) were generated using a 10x Chromium Controller (10x Genomics, Pleasanton, CA). GEM-RT was performed on the recovered GEMs in a C1000 Touch thermal cycler (Bio-Rad Laboratories, Hercules, CA) with the following program: 53°C for 45 min, 85°C for 5 min, and held at 4°C. After RT, single-stranded cDNA was recovered with Dynabeads MyOne Silane beads and amplified using a C1000 Touch thermal cycler. Amplified cDNA product was purified with HighPrep PCR Cleanup System beads using a 0.6x ratio (MagBio Genomics, Gaithersburg, MD; P/N AC-60050). cDNA was quantified using a Qubit instrument (Thermo Fisher Scientific, Waltham, MA) and visualized on a TapeStation D5000 tape (Agilent Technologies, Santa Clara, CA; P/N 5067-5588). cDNA was then converted to dual-indexed sequencing libraries by the following steps: enzymatic fragmentation, end-repair, and A-tailing, followed by double-sided bead size selection (0.6x and 0.8x ratios), adaptor ligation, post ligation bead clean up (0.8x ratio), indexing PCR with Dual Index Kit TT Set A (P/N 1000215) on a C1000 Touch thermal cycler. The final libraries were analyzed on a TapeStation 4200 system and quantified with Qubit and NEBNext library quantification kit (New England Biolabs, Ipswich, MA; P/N E7630). The libraries were then pooled at equimolar ratios and sequenced on Illumina NextSeq 500 sequencer (Illumina, San Diego, CA) with the following read lengths: read 128bp, i7 index 10bp, i5 index 10bp, and read 290bp. Simultaneously, the libraries were shipped to Element Biosciences for circularization and sequencing on their AVITI platform (Element Biosciences, San Diego, CA).

### snRNA sequencing data analysis

Single nuclei RNA sequencing files were converted into FASTQ and were aligned to the 10X prebuilt mouse reference, refdata-gex-mm10-2020-A with cellranger mkfastq. All libraries were visually QC checked using FastQC v0.11.7 and Fastq_screen v0.14.0 and confirmed to be of good quality. The gene expression matrices were generated using cellranger count function with default parameters, where reads with MAPQ < 255, intergenic reads, antisense reads, and reads mapping to >1 gene were removed from the dataset. CellRanger version 7.0.1 was used. All nuclei used for the following analysis contain no mitochondrial mRNAs. By this, the median genes per cell was 3568 for control and 3683 for Fmr1-cKO. Mean reads per cell was 65272 for control and 58232 for Fmr1-cKO. Seurat Version 5.0.1 and R Version 4.3.1 were used for the following analysis. The SCTransform function was run for normalization and regressing out variations. The RunPCA function was performed for the principal component analysis. The top 40 principal components were used to cluster the cells (FindNeighbors function, dim=40; FindClusters function, resolution=0.8). UMAP plot was generated by RunUMAP with dims=40. To extract differentially expressed genes (DEGs) in each cell type, the FindMarkers function was run. DEGs with p<0.02 in microglia and p<0.01 in PV neurons were subjected to the Ingenuity Pathway Analysis (IPA). Heatmaps were generated with ggplot2 version 3.4.4. Bar plots and volcano plot were produced by Prism version 10.2.0.

### FMRP immunoprecipitation

Male mice at P7-10 were used for immunoprecipitation. For each mouse, a half brain was used for immunoprecipitation, following a published protocol for FMRP immunoprecipitation^82^. Brains were homogenized in 1 ml ice-cold lysis buffer [10 mM HEPES (pH 7.4), 200 mM NaCl, 30 mM EDTA, 0.5% Triton X-100] with 1 U/μl SUPERase inhibitor, 1X protease inhibitor cocktail (Roche, 11873580001) and phosphatase inhibitor (PhosStop, Roche, 4906845001). Brain lysates were put on ice for 15 min and centrifuged for 10 min at 3000g at 4°C to remove nuclei. The supernatant was transferred to a new 2-ml round tube. NaCl concentration was raised to 400 mM, followed with centrifugation at 70,000g at 4°C for 20 min. Yeast tRNA at 100 μg/ml was added and the lysate was precleared with 35 μl protein G agarose (Roche, 11719416001) for 30 min at 4°C on a rotator. The lysate was centrifuged at 13,000g for 30s and supernatant was transferred to a new tube. For immunoprecipitation, 30 μl protein G agarose and 3 μl FMRP antibody (abcam, ab17722) were added to the supernatant and incubated at 4°C for 2 h on a rotator. The IP mixture was centrifuged at 13,000g for 30s to collect protein G agarose. Protein G agarose beads were washed 5 times with lysis buffer containing 0.1 U/ml SUPERase inhibitor and 10 μg/ml yeast tRNA (for the first four washes only). Bound RNAs were extracted with Arcturus PicoPure RNA isolation Kit (Applied biosystem, 12204-01) and cDNA was synthesized with qScript Ultra SuperMix (Quantabio). qPCR was done with SYBR Green master mix (Applied Biosystems, A25742). mRNA levels of candidate genes in the FMRP-immunoprecipitate were normalized to the β-actin level in the same immunoprecipitate. Enrichment of RNA was calculated as the ratio of the normalized mRNA level in WT precipitates to that in *Fmr1^-/y^* precipitates.

### Western blot

Protein extracts were prepared in RIPA buffer containing 50 mM Tris-HCl (pH7.4), 150 mM NaCl, 1% Triton X-100, 0.5% Sodium deoxylcholate, 1% SDS, 1 mM EDTA, 10 mM NaF, supplied with protease inhibitor cocktail (Roche, 11873580001) and phosphatase inhibitor (PhosStop, Roche, 4906845001). They were run in an SDS-PAGE gel and transferred onto a PVDF membrane. The membrane was blocked with 5% BSA in TBST for 1 h, followed with incubating with primary antibodies at 4°C overnight. Antibodies were diluted in 5% BSA in TBST. Primary antibodies were rabbit anti-FMRP (abcam, ab17722, 1:1000), mouse anti-α-Tub (Sigma, T6074, 1:10000), sheep anti-progranulin (R&D system, AF2557, 1:1000), rabbit anti-GAPDH (Cell signaling, 2118, 1:3000), rabbit anti-CTSD (Proteintech, 21327-1, 1:20000), mouse anti-HEXB (Santa Cruz, sc-376781, 1:25), and rabbit anti-Cystatin C (Proteintech, 12245-1-AP, 1:12000). IRDye infrared secondary antibodies (LI-COR Biosciences) were used: donkey anti-rabbit (#680LT) and donkey anti-mouse (#800CW). Secondary antibodies were diluted at 1:10,000 in blocking buffer and incubated for 1 h with the membrane. Odyssey Infrared Imaging System (Image Studio Lite Ver 4.0, LI-COR Biosciences) was used to detect the signals of target proteins. Alternatively, HRP-conjugated secondary antibodies were diluted in blocking buffer at 1:5000 and incubated with membranes for 1 h. Membranes were developed with SuperSignal West Pico PLUS Chemiluminescent Substrate (Thermo scientific, 34580) and imaged with Azure 600 gel imaging system. Protein levels were quantified using image J.

### Proximity ligation assay

Purified microglia were cultured in DMEM/F-12 + 10% FBS + 1X penicillin/streptomycin, on a coverslip pre-coated with 50 μg/ml Poly-D-lysine. On DIV2, microglia were treated with 3 μM puromycin (TOCRIS, 4089) in pre-warmed culture medium for 10 min. Cells were washed with pre-warmed DPBS twice, for 5 min each time. Cells were then fixed with 4% PFA for 15 min at room temperature. Cells were permeabilized by incubating with 0.3% Triton X-100 in PBS for 15 min. PLA was done with Sigma Duolink In Situ anti-rabbit PLUS probe (DUO92002) and anti-mouse MINUS probe (DUO92004). Procedures were performed according to the manufacturer’s manual. Briefly, cells were incubated with Duolink blocking buffer at 37°C for 60 min. Primary antibodies were diluted with Duolink antibody diluent and incubated with cells at 4°C overnight. Primary antibodies used were mouse anti-puromycin (DSHB, PMY-2A4, 1:100), rabbit anti-progranulin (Proteintech, 18410-1-AP, 1:2000), rabbit anti-CTSD (Proteintech, 21327-1-AP, 1:3000) and rabbit anti-GAPDH (Cell Signaling, #2118, 1:1200). Anti-rabbit PLUS probe and anti-mouse MINUS probe were diluted at 1:5 with antibody diluent and incubated with cells at 37°C for 60 min. Duolink In Situ detection reagent Red (Sigma, DUO92008) was used for PLA signal detection. Ligase were added to Duolink ligation buffer at a dilution of 1:40 and incubated with cells at 37°C for 30 min. Polymerase were diluted 1:80 with amplification buffer and incubated with cells at 37°C for 100 min. Prolong Gold mounting medium with DAPI was added and a coverslip was applied. Images were taken with Nikon C2 confocal microscope. Three male mice at P14 per genotype were used and 30 microglia were randomly selected for the quantification of PLA puncta with image J.

### Quantification of lysosomal volume and number in microglia

Fluorescent immunohistochemistry of Iba1 and CD68 were performed with IC tissues from male mice at P14. Four z-stack confocal images per mouse, at a thickness of 15 μm with a 0.25-μm interval, were acquired with a 60X oil lens using a Nikon C2 confocal microscope. CD68 volume and Iba1 volume for each microglia were measured using ‘surface’ function in Imaris software 10.0.0 (Oxford Instruments).

### Quantification of cells undergoing apoptosis in the IC

Every other sections of IC tissues from male mice at P14 were used in triple immunohistochemistry for cleaved casp3, PV and Iba1. The sections were screened under Nikon C2 confocal microscope with 40X lens to count the number of cells positive for cleaved caspase-3. Once encountering a casp3^+^ cell, co-expression with PV or Iba1 was checked under the confocal microscopy.

### Quantification of pTDP43 puncta

Every other IC sections from P14 mice were used for immunostaining of pTDP43, PV and Iba1. Sections were examined under Nikon confocal microscope with a 60X oil lens. To quantify pTDP43 puncta density, 14-17 images per mouse were randomly taken in the IC. pTDP43 puncta number and size were quantified with image J. To quantify pTDP43 puncta associated with apoptosis, IC sections were screened under the microscope. Cells with ongoing apoptosis were identified by the presence of a fragmented nucleus.

### Quantification of Lamp1 puncta number and size in PV neurons

Every other sections of the IC from P14 mice were used for co-staining of PV and Lamp1. Ten images were randomly acquired for each mouse with a 60X oil lens under a Nikon C2 confocal microscope. Density and size of Lamp1 puncta in the IC were quantified with image J. Ten PV neurons were randomly selected per mouse, and Lamp1 puncta inside the soma of each PV neuron were analyzed by image J.

### Assays for lysosomal functionality

Purified microglia were cultured in DMEM/F-12 + 10% FBS + 1X penicillin/streptomycin. The assays were performed following published methods with minor optimization^83^. For quantification of lysosome acidity, cells after 24 h in culture were treated with 1 μM Lysosensor Green DND-189 (Invitrogen L7535) for 20 min in a 37°C incubator. Images were taken with Nikon C2 confocal microscope, and mean green fluorescence in each cell was quantified with image J. In order to quantify lysosomal degradation capability in microglia, microglia on DIV1 were treated with 10 μg/ml DQ-BSA (Invitrogen D12050) for 16 h in a 37°C incubator. Cells were washed with prewarmed DPBS twice for 5 min each time. The cells were then fixed with 4% PFA for 15 min at room temperature. Immunocytochemistry to probe Iba1 (Wako, 019-19741, 1:500) and Lamp1 (DSHB, 1D4B, 1:250) was perform. Confocal images were acquired, and DQ-BSA fluorescence in each Iba1^+^ cell was quantified with image J. Meanwhile, Zeiss Elyra SIM super-resolution microscope was used to show that DQ-BSA puncta in Iba1^+^ cells were colocalized with Lamp1^+^ lysosomes. Images were taken with a 63X oil lens.

### Viral expression of progranulin in the brain of *Fmr1*^-/y^ null mice

Plasmid pAAV-CAG-mNeonGreen was purchased from addgene (#99134). Plasmid pAAV-CAG-Grn was generated by replacing the mNeonGreen cassette with the *Grn* coding sequence. To validate the functionality of the two plasmid, HEK293T cells were transfected with the two plasmids separately. 48h after transfection, mNeonGreen expression was confirmed under a fluorescence microscope. Cells were lysed, and progranulin overexpression was examined with Western blot. After the validation, AAV8-CAG-mNeonGreen and AAV8-CAG-Grn were packaged and purified using TaKaRa AAVpro Purification Kit Maxi (Cat. #6666) according to the product manual. Briefly, HEK293T cells were grown in 150-mm culture plates. When the culture achieved 80% of confluency, cells were transfected with a DNA mixture containing 3 plasmids: pAAV-CAG-mNeonGreen or pAAV-CAG-Grn, pAAV2/8 (addgene plasmid #112864) and pAdDeltaF6 (addgene plasmid # 112867). Viruses were collected 96 h after transfection and purified. The titer of the virus preparation was determined using qPCR. To rescue pathological and behavioral deficits in *Fmr1^-/y^*null mice, AAV8-CAG-mNeonnGreen and AAV8-CAG-Grn were diluted by 10 times with PBS. 1.5 μl of diluted viruses were intracerebroventricularly injected into the lateral ventricle of each hemisphere of the brain at P0. Brains were collected at P22 for measurement of PV neuron density in the inferior colliculus and somatosensory cortex, as described earlier. Alternatively, mice at P22-25 were used for test of audiogenic seizures.

### Statistics

All data in figures are presented as mean ± SEM. Statistical methods used are described in figure legends, and statistical analysis was done using GraphPad Prism Version 10.2.0. Cell counting was done by an examiner blind to the genotypes of the mice.

## Acknowledgements

We thank Dr. David Nelson at Baylor College of Medicine for kindly providing us with the *Fmr1^lox^* mouse strain, Dr. Ronald Davis at UF Scripps for comments on the manuscript, Dr. Robert Witwicki at the UF Scripps Genomic Sequencing Core for snRNA sequencing, and Alexander Trouern-Trend at the UF Scripps Bioinformatics and Statistics Core for assistance in analyzing snRNA-Seq data. This work was supported by grants from the U. S. National Institutes of Health to B.X. (R01 MH125187) and E.J.H. (R01 AG081452 and R01 AG057462). E.J.H. is an investigator with the Bluefield Project to Cure FTD. H.L. was partially supported by a Training Grant in Alzheimer’s Drug Discovery from the Lottie French Lewis Fund of the Community Foundation for Palm Beach and Martin Counties.

## Author Contributions

H.L. and B.X. conceived and designed the experiments and interpreted the data. E.J.H. provided insights in some experimental designs. H.L. performed all experiments and analyzed all data. J.J.Y.C. contributed to analysis of snRNA-Seq data. H.L. and B.X. wrote the paper with comments from E.J.H.

## References

1. Hagerman, R.J., et al. Fragile X syndrome. Nat Rev Dis Primers 3, 17065 (2017).

2. Bagni, C. & Zukin, R.S. A Synaptic Perspective of Fragile X Syndrome and Autism Spectrum Disorders. Neuron 101, 1070–1088 (2019).

3. Richter, J.D. & Zhao, X. The molecular biology of FMRP: new insights into fragile X syndrome. Nat Rev Neurosci (2021).

4. Nelson, D.L., Orr, H.T. & Warren, S.T. The unstable repeats--three evolving faces of neurological disease. Neuron 77, 825–843 (2013).

5. Wang, L.W., Berry-Kravis, E. & Hagerman, R.J. Fragile X: leading the way for targeted treatments in autism. Neurotherapeutics 7, 264–274 (2010).

6. Clifford, S., et al. Autism spectrum phenotype in males and females with fragile X full mutation and premutation. J Autism Dev Disord 37, 738–747 (2007).

7. Harris, S.W., et al. Autism profiles of males with fragile X syndrome. Am J Ment Retard 113, 427–438 (2008).

8. Pieretti, M., et al. Absence of expression of the FMR-1 gene in fragile X syndrome. Cell 66, 817–822 (1991).

9. Verkerk, A.J., et al. Identification of a gene (FMR-1) containing a CGG repeat coincident with a breakpoint cluster region exhibiting length variation in fragile X syndrome. Cell 65, 905–914 (1991).

10. Brown, V., et al. Microarray identification of FMRP-associated brain mRNAs and altered mRNA translational profiles in fragile X syndrome. Cell 107, 477–487 (2001).

11. Darnell, J.C., et al. FMRP stalls ribosomal translocation on mRNAs linked to synaptic function and autism. Cell 146, 247–261 (2011).

12. Zalfa, F., et al. The fragile X syndrome protein FMRP associates with BC1 RNA and regulates the translation of specific mRNAs at synapses. Cell 112, 317–327 (2003).

13. Lu, R., et al. The fragile X protein controls microtubule-associated protein 1B translation and microtubule stability in brain neuron development. Proc Natl Acad Sci U S A 101, 15201–15206 (2004).

14. Osterweil, E.K., Krueger, D.D., Reinhold, K. & Bear, M.F. Hypersensitivity to mGluR5 and ERK1/2 leads to excessive protein synthesis in the hippocampus of a mouse model of fragile X syndrome. J Neurosci 30, 15616–15627 (2010).

15. Gross, C., et al. Excess phosphoinositide 3-kinase subunit synthesis and activity as a novel therapeutic target in fragile X syndrome. J Neurosci 30, 10624–10638 (2010).

16. Napoli, I., et al. The fragile X syndrome protein represses activity-dependent translation through CYFIP1, a new 4E-BP. Cell 134, 1042–1054 (2008).

17. Nishimura, Y., et al. Genome-wide expression profiling of lymphoblastoid cell lines distinguishes different forms of autism and reveals shared pathways. Hum Mol Genet 16, 1682–1698 (2007).

18. Bechara, E.G., et al. A novel function for fragile X mental retardation protein in translational activation. PLoS Biol 7, e16 (2009).

19. Gross, C., Yao, X., Pong, D.L., Jeromin, A. & Bassell, G.J. Fragile X mental retardation protein regulates protein expression and mRNA translation of the potassium channel Kv4.2. J Neurosci 31, 5693–5698 (2011).

20. Zalfa, F., et al. A new function for the fragile X mental retardation protein in regulation of PSD-95 mRNA stability. Nat Neurosci 10, 578–587 (2007).

21. Zhang, F., et al. Fragile X mental retardation protein modulates the stability of its m6A-marked messenger RNA targets. Hum Mol Genet 27, 3936–3950 (2018).

22. McMahon, A.C., et al. TRIBE: Hijacking an RNA-Editing Enzyme to Identify Cell-Specific Targets of RNA-Binding Proteins. Cell 165, 742–753 (2016).

23. Tran, S.S., et al. Widespread RNA editing dysregulation in brains from autistic individuals. Nat Neurosci 22, 25–36 (2019).

24. Contractor, A., Ethell, I.M. & Portera-Cailliau, C. Cortical interneurons in autism. Nat Neurosci 24, 1648–1659 (2021).

25. Selby, L., Zhang, C. & Sun, Q.Q. Major defects in neocortical GABAergic inhibitory circuits in mice lacking the fragile X mental retardation protein. Neurosci Lett 412, 227–232 (2007).

26. Kourdougli, N., et al. Improvement of sensory deficits in fragile X mice by increasing cortical interneuron activity after the critical period. Neuron 111, 2863–2880 e2866 (2023).

27. Juarez, P., Salcedo-Arellano, M.J., Dufour, B. & Martinez-Cerdeno, V. Fragile X cortex is characterized by decreased parvalbumin-expressing interneurons. Cereb Cortex 34(2024).

28. Hashemi, E., Ariza, J., Rogers, H., Noctor, S.C. & Martinez-Cerdeno, V. The Number of Parvalbumin-Expressing Interneurons Is Decreased in the Medial Prefrontal Cortex in Autism. Cereb Cortex 27, 1931–1943 (2017).

29. Ariza, J., Rogers, H., Hashemi, E., Noctor, S.C. & Martinez-Cerdeno, V. The Number of Chandelier and Basket Cells Are Differentially Decreased in Prefrontal Cortex in Autism. Cereb Cortex 28, 411–420 (2018).

30. Gogolla, N., et al. Common circuit defect of excitatory-inhibitory balance in mouse models of autism. J Neurodev Disord 1, 172–181 (2009).

31. Canetta, S., et al. Maternal immune activation leads to selective functional deficits in offspring parvalbumin interneurons. Mol Psychiatry 21, 956–968 (2016).

32. Yu, D., et al. Microglial GPR56 is the molecular target of maternal immune activation-induced parvalbumin-positive interneuron deficits. Sci Adv 8, eabm2545 (2022).

33. Shin Yim, Y., et al. Reversing behavioural abnormalities in mice exposed to maternal inflammation. Nature 549, 482–487 (2017).

34. Godavarthi, S.K., Sharma, A. & Jana, N.R. Reversal of reduced parvalbumin neurons in hippocampus and amygdala of Angelman syndrome model mice by chronic treatment of fluoxetine. J Neurochem 130, 444–454 (2014).

35. Chen, Z., et al. Accumulated quiescent neural stem cells in adult hippocampus of the mouse model for the MECP2 duplication syndrome. Scientific reports 7, 41701 (2017).

36. Li, X.G., Somogyi, P., Tepper, J.M. & Buzsaki, G. Axonal and dendritic arborization of an intracellularly labeled chandelier cell in the CA1 region of rat hippocampus. Exp Brain Res 90, 519–525 (1992).

37. Marin, O. Developmental timing and critical windows for the treatment of psychiatric disorders. Nat Med 22, 1229–1238 (2016).

38. Thion, M.S. & Garel, S. On place and time: microglia in embryonic and perinatal brain development. Curr Opin Neurobiol 47, 121–130 (2017).

39. Salter, M.W. & Stevens, B. Microglia emerge as central players in brain disease. Nat Med 23, 1018–1027 (2017).

40. Peri, F. & Nusslein-Volhard, C. Live imaging of neuronal degradation by microglia reveals a role for v0-ATPase a1 in phagosomal fusion in vivo. Cell 133, 916–927 (2008).

41. Marin-Teva, J.L., et al. Microglia promote the death of developing Purkinje cells. Neuron 41, 535–547 (2004).

42. Sedel, F., Bechade, C., Vyas, S. & Triller, A. Macrophage-derived tumor necrosis factor alpha, an early developmental signal for motoneuron death. J Neurosci 24, 2236–2246 (2004).

43. Mazaheri, F., et al. Distinct roles for BAI1 and TIM-4 in the engulfment of dying neurons by microglia. Nat Commun 5, 4046 (2014).

44. Tremblay, M.E., Lowery, R.L. & Majewska, A.K. Microglial interactions with synapses are modulated by visual experience. PLoS Biol 8, e1000527 (2010).

45. Wake, H., Moorhouse, A.J., Miyamoto, A. & Nabekura, J. Microglia: actively surveying and shaping neuronal circuit structure and function. Trends Neurosci 36, 209–217 (2013).

46. Stevens, B., et al. The classical complement cascade mediates CNS synapse elimination. Cell 131, 1164–1178 (2007).

47. Schafer, D.P., et al. Microglia sculpt postnatal neural circuits in an activity and complement-dependent manner. Neuron 74, 691–705 (2012).

48. Tanaka, Y., et al. Progranulin regulates lysosomal function and biogenesis through acidification of lysosomes. Hum Mol Genet 26, 969–988 (2017).

49. Zhou, X., et al. Impaired prosaposin lysosomal trafficking in frontotemporal lobar degeneration due to progranulin mutations. Nat Commun 8, 15277 (2017).

50. Lui, H., et al. Progranulin Deficiency Promotes Circuit-Specific Synaptic Pruning by Microglia via Complement Activation. Cell 165, 921–935 (2016).

51. Kao, A.W., McKay, A., Singh, P.P., Brunet, A. & Huang, E.J. Progranulin, lysosomal regulation and neurodegenerative disease. Nat Rev Neurosci 18, 325–333 (2017).

52. Zhang, J., et al. Neurotoxic microglia promote TDP-43 proteinopathy in progranulin deficiency. Nature 588, 459–465 (2020).

53. Gholizadeh, S., Halder, S.K. & Hampson, D.R. Expression of fragile X mental retardation protein in neurons and glia of the developing and adult mouse brain. Brain Res 1596, 22–30 (2015).

54. Koekkoek, S.K., et al. Deletion of FMR1 in Purkinje cells enhances parallel fiber LTD, enlarges spines, and attenuates cerebellar eyelid conditioning in Fragile X syndrome. Neuron 47, 339–352 (2005).

55. Parkhurst, C.N., et al. Microglia promote learning-dependent synapse formation through brain-derived neurotrophic factor. Cell 155, 1596–1609 (2013).

56. Madisen, L., et al. A robust and high-throughput Cre reporting and characterization system for the whole mouse brain. Nat Neurosci 13, 133–140 (2010).

57. Louros, S.R., et al. Excessive proteostasis contributes to pathology in fragile X syndrome. Neuron 111, 508–525 e507 (2023).

58. Musumeci, S.A., et al. Audiogenic seizure susceptibility is reduced in fragile X knockout mice after introduction of FMR1 transgenes. Exp Neurol 203, 233–240 (2007).

59. Gonzalez, D., et al. Audiogenic Seizures in the Fmr1 Knock-Out Mouse Are Induced by Fmr1 Deletion in Subcortical, VGlut2-Expressing Excitatory Neurons and Require Deletion in the Inferior Colliculus. J Neurosci 39, 9852–9863 (2019).

60. Xu, Z.X., et al. Elevated protein synthesis in microglia causes autism-like synaptic and behavioral aberrations. Nat Commun 11, 1797 (2020).

61. Wang, A., et al. Innate immune sensing of lysosomal dysfunction drives multiple lysosomal storage disorders. Nat Cell Biol 26, 219–234 (2024).

62. Di, Y.Q., et al. Autophagy triggers CTSD (cathepsin D) maturation and localization inside cells to promote apoptosis. Autophagy 17, 1170–1192 (2021).

63. Fernandez-Mosquera, L., et al. Mitochondrial respiratory chain deficiency inhibits lysosomal hydrolysis. Autophagy 15, 1572–1591 (2019).

64. Asrani, K., et al. mTORC1 feedback to AKT modulates lysosomal biogenesis through MiT/TFE regulation. J Clin Invest 129, 5584–5599 (2019).

65. Goswami, A.B., Karadarevic, D. & Castano-Rodriguez, N. Immunity-related GTPase IRGM at the intersection of autophagy, inflammation, and tumorigenesis. Inflamm Res 71, 785–795 (2022).

66. Maurin, T., et al. HITS-CLIP in various brain areas reveals new targets and new modalities of RNA binding by fragile X mental retardation protein. Nucleic Acids Res 46, 6344–6355 (2018).

67. Ascano, M., Jr., et al. FMRP targets distinct mRNA sequence elements to regulate protein expression. Nature 492, 382–386 (2012).

68. Life, B., Bettio, L.E.B., Gantois, I., Christie, B.R. & Leavitt, B.R. Progranulin is an FMRP target that influences macroorchidism but not behaviour in a mouse model of Fragile X Syndrome. Curr Res Neurobiol 5, 100094 (2023).

69. Xu, H. & Ren, D. Lysosomal physiology. Annu Rev Physiol 77, 57–80 (2015).

70. Kiselyov, K., Jennigs, J.J., Jr., Rbaibi, Y. & Chu, C.T. Autophagy, mitochondria and cell death in lysosomal storage diseases. Autophagy 3, 259–262 (2007).

71. Rhinn, H., Tatton, N., McCaughey, S., Kurnellas, M. & Rosenthal, A. Progranulin as a therapeutic target in neurodegenerative diseases. Trends Pharmacol Sci 43, 641–652 (2022).

72. Plotegher, N. & Duchen, M.R. Mitochondrial Dysfunction and Neurodegeneration in Lysosomal Storage Disorders. Trends Mol Med 23, 116–134 (2017).

73. Baker, M., et al. Mutations in progranulin cause tau-negative frontotemporal dementia linked to chromosome 17. Nature 442, 916–919 (2006).

74. Li, M., et al. Identification of FMR1-regulated molecular networks in human neurodevelopment. Genome Res 30, 361–374 (2020).

75. Dolen, G., et al. Correction of fragile X syndrome in mice. Neuron 56, 955–962 (2007).

76. Kerr, J.S., et al. Mitophagy and Alzheimer’s Disease: Cellular and Molecular Mechanisms. Trends Neurosci 40, 151–166 (2017).

77. Ruden, J.B., Dugan, L.L. & Konradi, C. Parvalbumin interneuron vulnerability and brain disorders. Neuropsychopharmacology 46, 279–287 (2021).

78. Mientjes, E.J., et al. The generation of a conditional Fmr1 knock out mouse model to study Fmrp function in vivo. Neurobiol Dis 21, 549–555 (2006).

79. Osterweil, E.K., et al. Lovastatin corrects excess protein synthesis and prevents epileptogenesis in a mouse model of fragile X syndrome. Neuron 77, 243–250 (2013).

80. Bohlen, C.J., et al. Diverse Requirements for Microglial Survival, Specification, and Function Revealed by Defined-Medium Cultures. Neuron 94, 759–773 e758 (2017).

81. Stogsdill, J.A., et al. Pyramidal neuron subtype diversity governs microglia states in the neocortex. Nature 608, 750–756 (2022).

82. Edbauer, D., et al. Regulation of synaptic structure and function by FMRP-associated microRNAs miR-125b and miR-132. Neuron 65, 373–384 (2010).

83. Wang, B., et al. TFEB-vacuolar ATPase signaling regulates lysosomal function and microglial activation in tauopathy. bioRxiv (2023).

