## Supplemental Figures for "Loss of FMRP in microglia promotes degeneration of parvalbumin neurons and audiogenic seizures via progranulin insufficiency"

**Extended Data Fig. 1: FMRP expression in microglia of postnatal brains.**

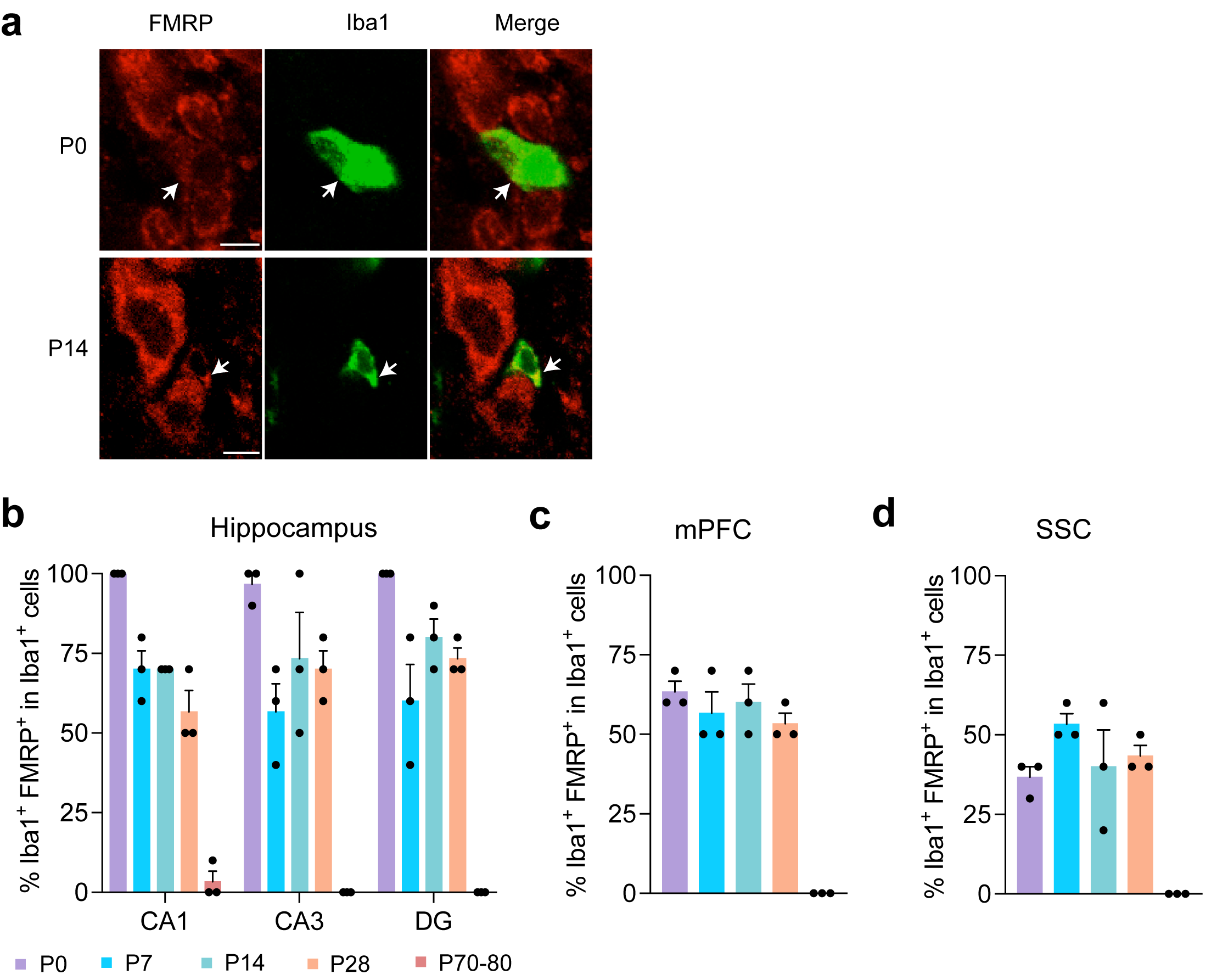

**a**, Immunohistochemistry of FMRP and Iba1 in the IC of WT mice at P0 and P14. White arrows indicate colocalization. Scale bars, 5 µm. **b-d**, Percentage of microglia with FMRP expression in the hippocampus (**b**), medial prefrontal cortex (mPFC, **c**), and somatosensory cortex (SSC, **d**) in mice at P0, P7, P14, P28, and P70-80. n=3 mice of both sexes for each age.

**Extended Data Fig. 2: Selective deletion of the *Fmr1* gene in microglia leads to audiogenic seizures.**

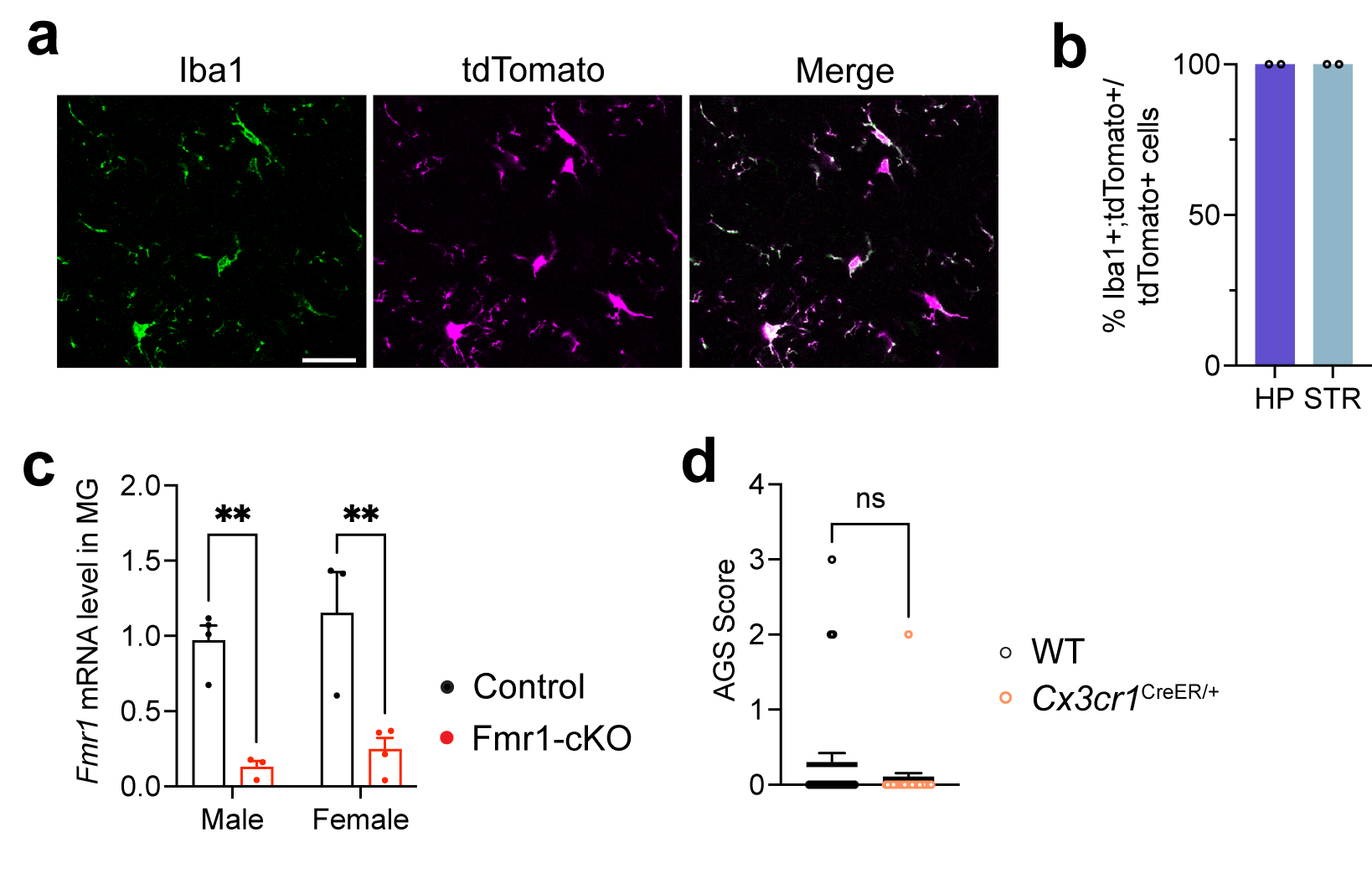

**a**, Confocal images showing complete overlap of tdTomato-expressing cells with Iba1-expressing cells in *Cx3cr1^CreER/+^*;Ai9/+ mice. This suggests that the Cre recombinase activity in the *Cx3cr1^CreER/+^* mouse strain is restricted to microglia. Scale bar: 30 μm. **b**, Percentage of tdTomato^+^ cells that are positive for Iba1 reactivity. 100-150 tdTomato^+^ cells were checked in each brain region for each mouse. Two Cx3cr1^CreER/+^;Ai9/+ mice at P14 were examined. HP, hippocampus; STR, striatum. **c**, The *Fmr1* mRNA level was decreased in microglia isolated from either male or female Fmr1 cKO mice. Two-way ANOVA followed with Bonferroni’s multiple comparisons test, **p<0.01, n=3-4 mice per group. **d**, Distribution of AGS scores in Cx3cr1^CreER/+^ mice and WT littermates. Mann-Whitney test, ns: not significant, n=26 mice per genotype, including both males and females.

**Extended Data Fig. 3: Microglial *Fmr1* deletion leads to changes in microglia and parvalbumin neurons in the IC.**

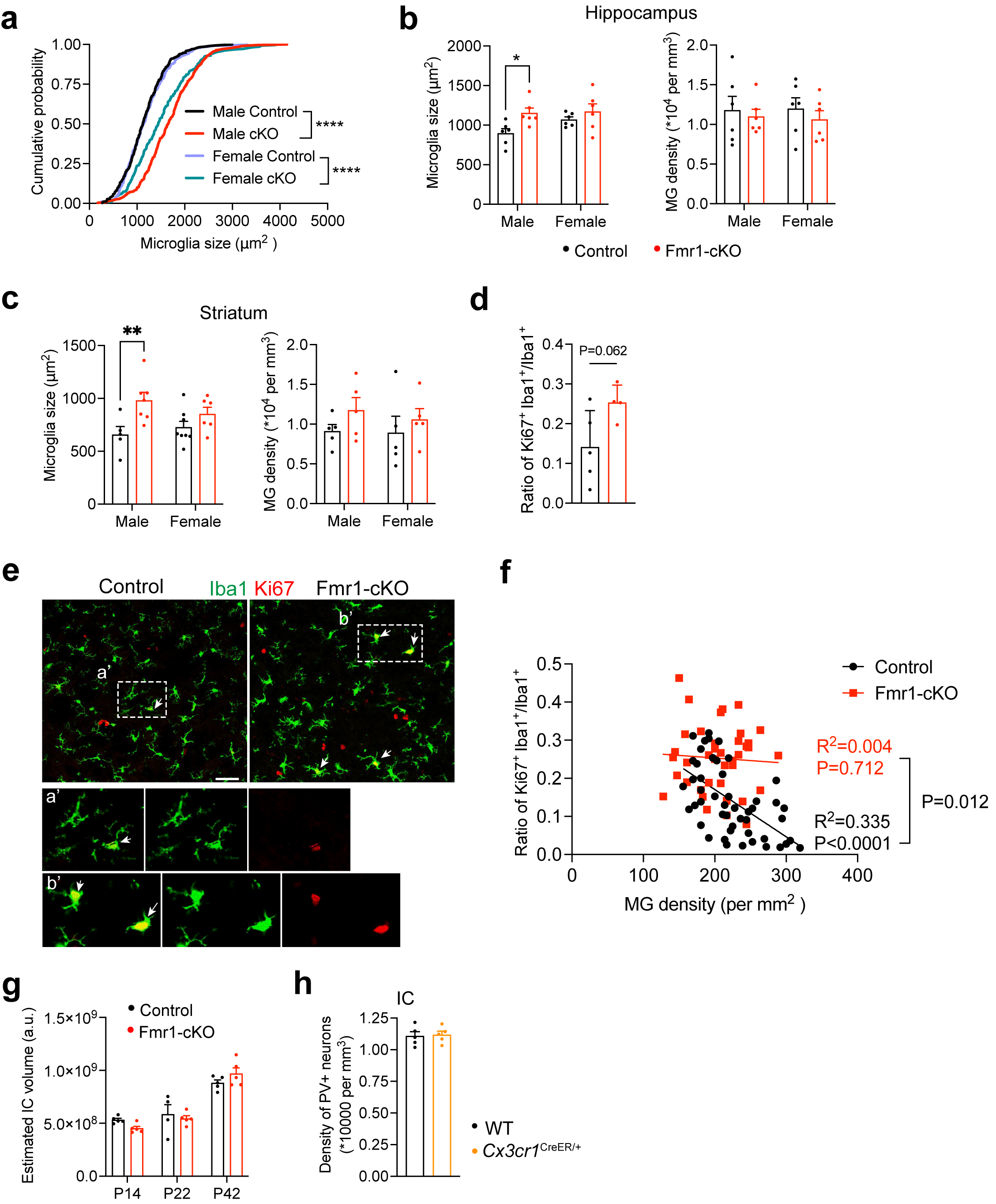

**a**, Cumulative probability of microglial size in the IC of Control and Fmr1-cKO mice at P14. Kolmogorov-Smirnov test, ****p<0.0001, n=5 mice per group and 50-60 microglia from each mouse. **b**, Size and density of microglia in the hippocampus of Control and Fmr1-cKO mice at P14. Two-way ANOVA followed with Bonferroni’s multiple comparisons test, *p<0.05, n=6 mice per group. **c**, Size and density of microglia in the striatum of Control and Fmr1-cKO mice at P14. Two-way ANOVA followed with Bonferroni’s multiple comparisons test, **p<0.01, n=6 mice per group. **d**, Ratio of proliferating microglia over all microglia, showing a trend of increase in the IC of male Fmr1-cKO mice. Unpaired t test, n=4-5 mice per group. **e**, Representative confocal images showing colocalization of Iba1 and Ki67 in the IC of male Control and Fmr1-cKO mice at P10. Arrows indicates colocalization. Scale bar, 50 μm. The areas within the dotted squares were enlarged in the bottom images. **f**, Linear regression between microglia density and ratio of proliferating microglia over all microglia in the IC of Control and Fmr1-cKO mice at P10. n=4-5 mice per group. **g**, Volume of the IC in one hemisphere is normal in Fmr1-cKO mice at different ages. Two-way ANOVA, n=4-5 mice per group. h, Density of PV neurons is normal in the IC of male Cx3cr1^CreER/+^ mice at P22. Two-tailed unpaired t test, p=0.83, n=5 mice per group.

Extended Data **Fig. 4: snRNA sequencing of IC tissues dissected from Control and Fmr1-cKO mice at P22**.

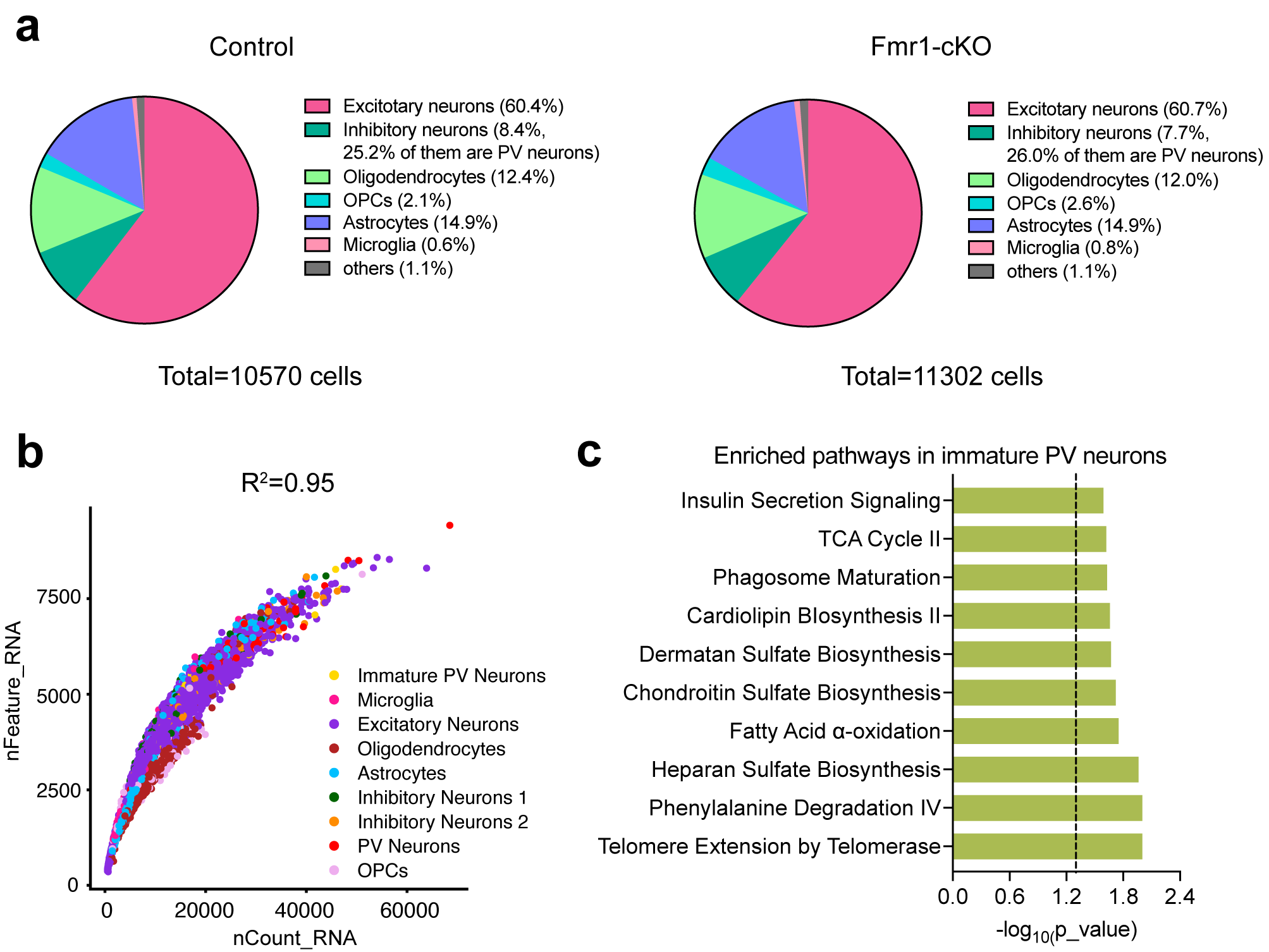

**a**, Cell composition of the snRNA-seq data from the IC of control and Fmr1-cKO mice. **b**, QC plot showing the high correlation (R^2^=0.95) between the number of RNA reads (nCount_RNA) and the number of detected genes (nFeature_RNA) in each cell. Cell types are color-coded. **c**, Top 10 enriched signaling pathways in immature PV neurons of Fmr1-cKO mice. Dashed line indicates p=0.05.

Extended Data **Fig. 5: Microglia with FMRP depletion displayed lysosomal dysfunction.**

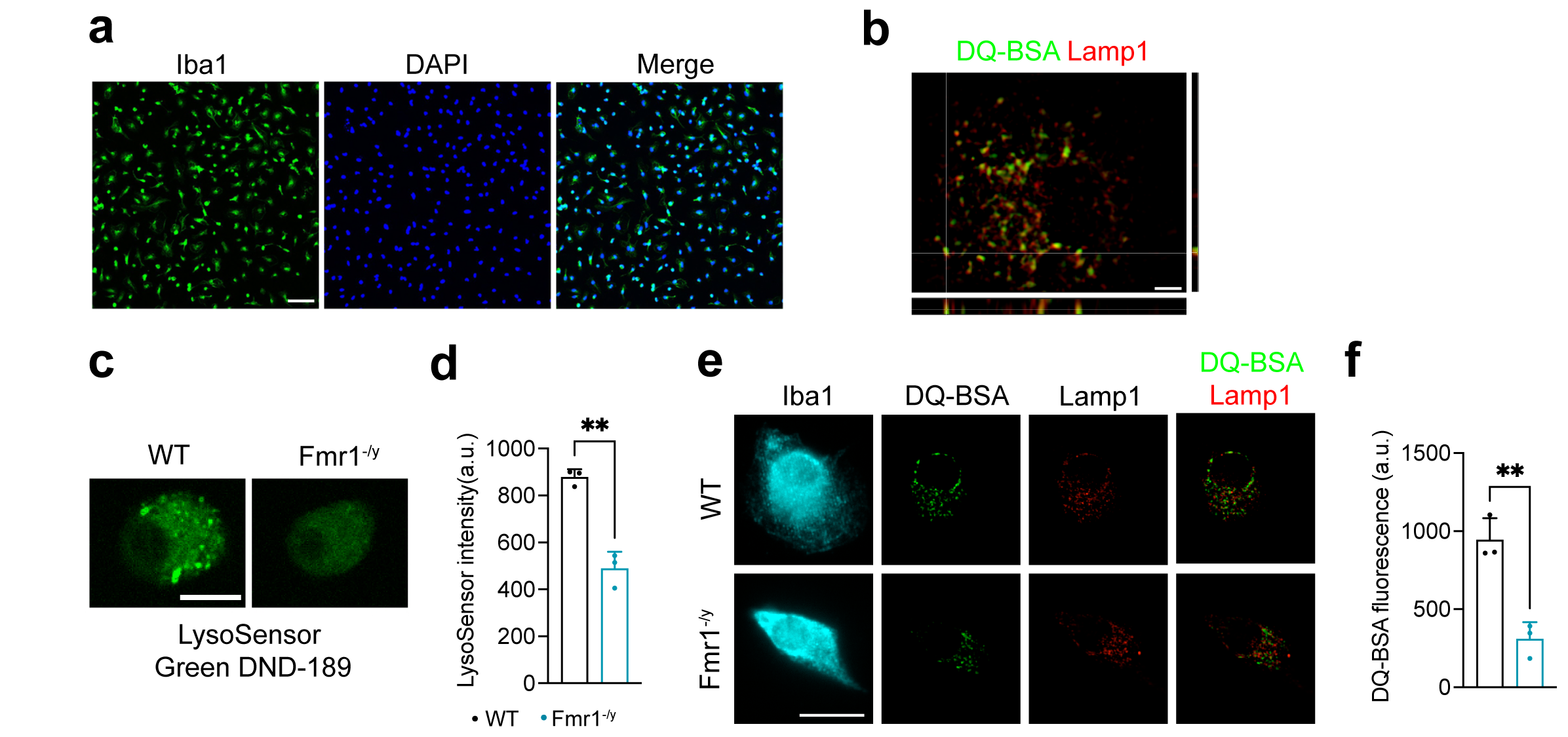

**a**, Confocal images showing Iba1 immunoreactivity in primary microglia purified from P14 mice. The vast majority of the purified cells were Iba1^+^ microglia. Scale bar, 50 μm. **b**, Z stack images were taken with a super-resolution structured illumination microscope, and 3D was reconstructed with Imaris software. Orthogonal images were generated to show the colocalization of DQ-BSA and Lamp1 in microglia from control mice. scale bar: 2 μm. **c**, Representative confocal images showing the LysoSensor Green DND-189 fluorescence in microglia purified from WT and *Fmr1*^-/y^ mice at P12-P14. Scale bar: 10 μm. **d**, LysoSensor fluorescence intensity were decreased in microglia from *Fmr1*^-/y^ mice compared with those from male WT mice. Unpaired t test, **p<0.01, n=3 mice per genotype, 10-20 microglia were quantified per mouse. **e**, Representative images taken with a super-resolution microscope showing colocalization of DQ-BSA (green) and Lamp1(red) puncta in Iba1^+^ microglia purified from male WT and *Fmr1*^-/y^ mice. Scale bar: 10 μm. **f**, DQ-BSA fluorescence was decreased in microglia purified from *Fmr1*^-/y^ mice. Unpaired t test, **p<0.01, n=3 mice per genotype, 10-20 microglia were quantified per mouse.

Extended Data **Fig. 6: Lysosomal dysfunction of PV neurons in Fmr1-cKO mice**.

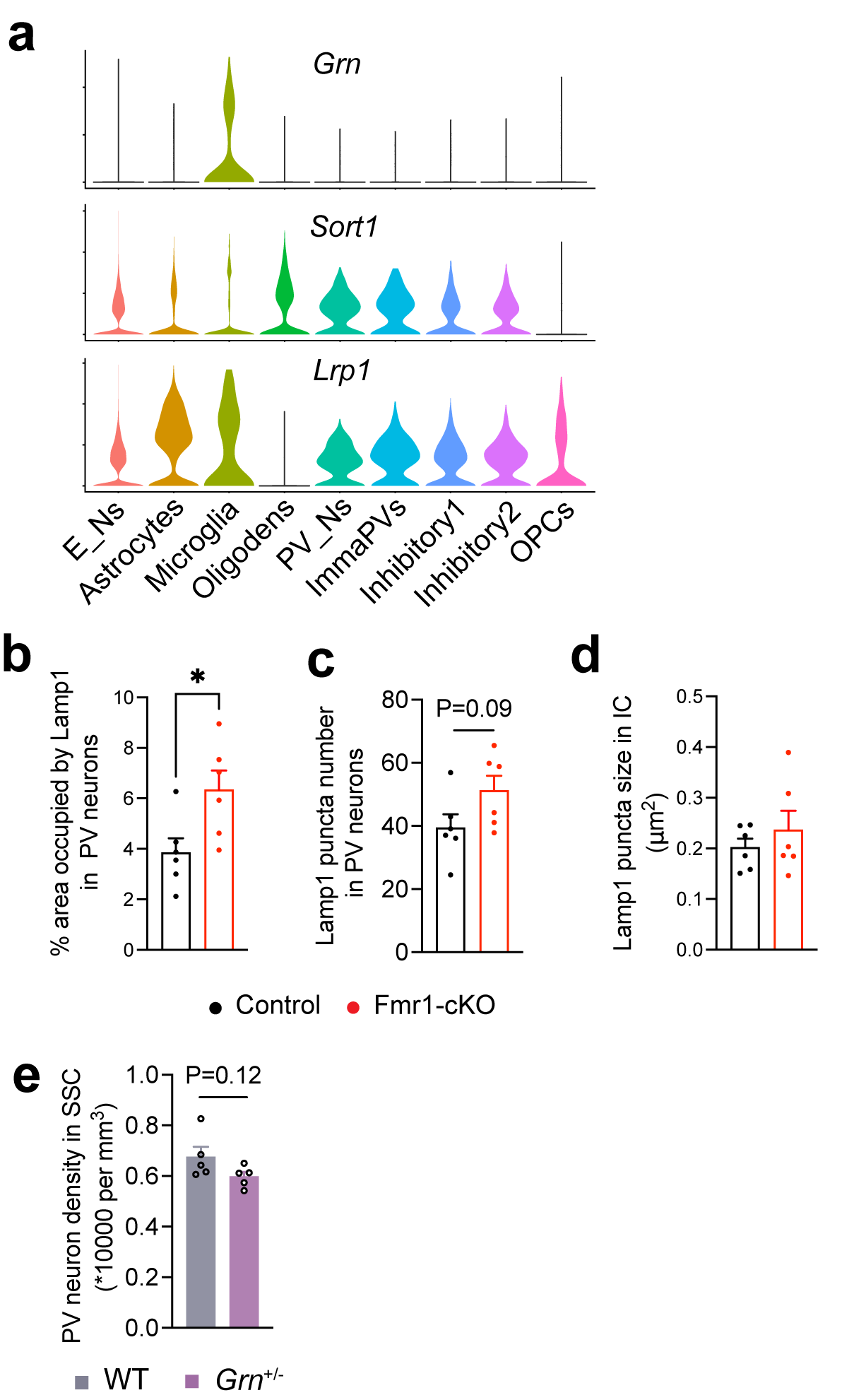

**a**, Violin plots showing the expression of *Grn* and its receptors *Sort1* and *Lrp1* in IC of P22 mice. Cell types were color-coded. The plots were based on the snRNA-seq data showing in Fig. 4. E_Ns, Excitatory neurons; Oligodens, Oligodendrocytes; PV_Ns, PV neurons; ImmaPVs: Immature PV neurons. **b**, Percentage of PV neuron area occupied by Lamp1 immunoreactivity in the IC of male Control and Fmr1-cKO mice. unpaired t test, *p<0.05, n=6 male mice per genotype. **c**, There is a trend of increase in the number of Lamp1^+^ puncta in PV neurons of Fmr1-cKO mice. Unpaired t test, n=6 male nice per genotype. **d**, Lamp1 puncta size was not changed across the whole IC section in Fmr1-cKO mice. n=6 male mice per genotype. **e**, Density of PV neurons in SSC of WT and *Grn*^+/-^ mice at P22. Unpaired t test, n=5 male mice per genotype.

Extended Data **Fig. 7:** **Viral PGRN overexpression in the brain of *Fmr1* KO mice.**

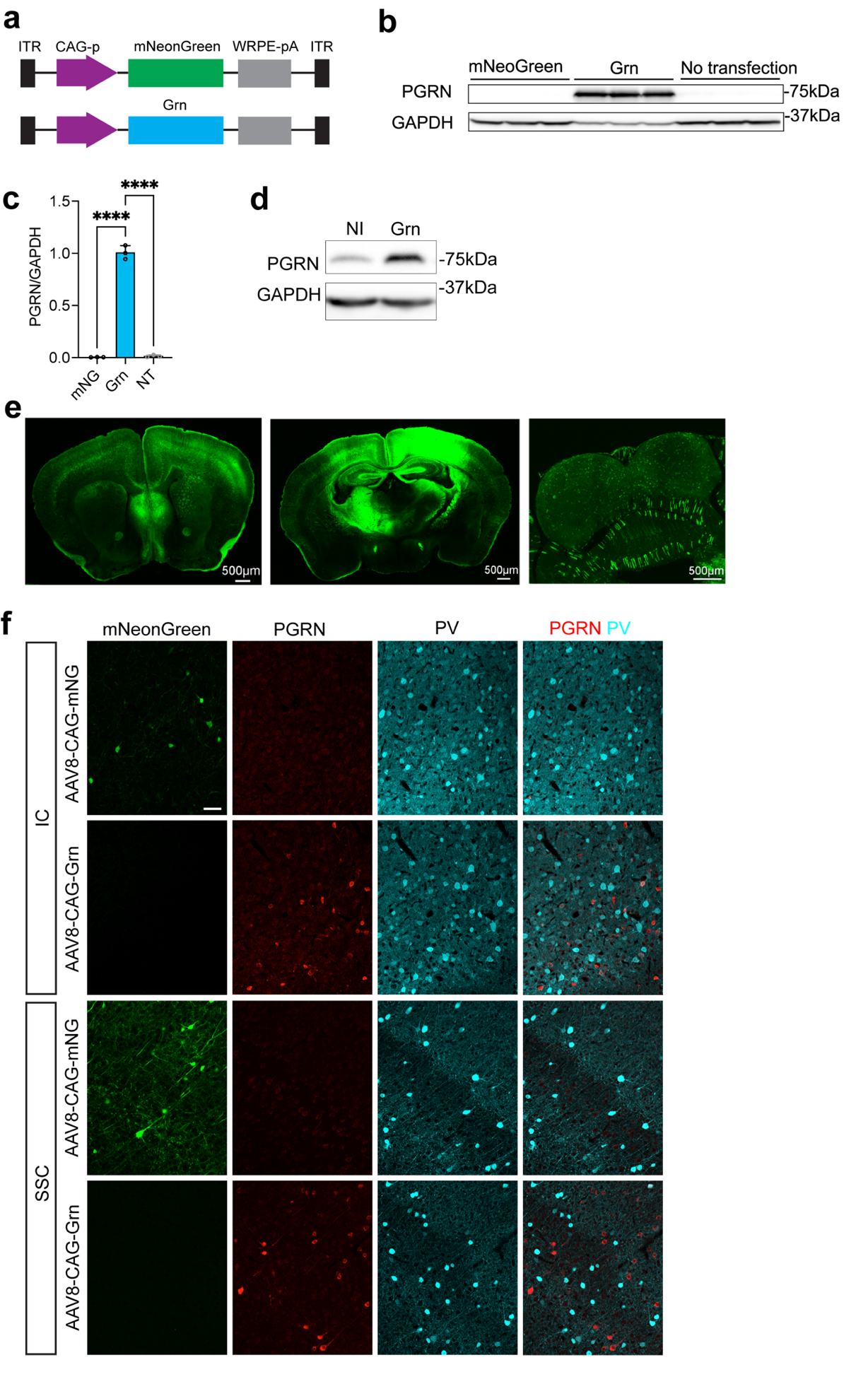

a, Diagram showing the structure of viral vectors expressing mNeonGreen or Grn. **b**, Western blots showing PGRN expression in HEK293T cells after transfection. **c**, Quantification of PGRN levels in HEK293T cells transfected with the construct expressing mNeonGreen (mNG) or Grn or without transfection (NT). One-way ANOVA followed with Tukey’s multiple comparisons test, ****p<0.0001, n=3. **d**, Western blots showing PGRN expression in the IC of WT mice at P14 after intracerebroventricular injection of AAV8-CAG-Grn (Grn) at P0 or no injection (NI). **e**, Representative confocal images showing the mNeonGreen expression throughout the brain at P21. AAV8-CAG-mNeonGreen was bilaterally injected into the lateral ventricle of the brain at P0. **f**, Representative confocal images showing mNeonGreen and PGRN expression in the IC and SSC of mice at P22 after intracerebroventricular injection of viruses at P0. Scale bar, 50 μm.
